# Structures of protein folding intermediates on the ribosome

**DOI:** 10.1101/2025.04.07.647236

**Authors:** Sammy H.S. Chan, Julian O. Streit, Tomasz Włodarski, Alkistis N. Mitropoulou, Anaïs M.E. Cassaignau, Ivana V. Bukvin, Lisa D. Cabrita, John Christodoulou

## Abstract

The ribosome biases the conformations sampled by nascent polypeptide chains along folding pathways towards biologically active states. A hallmark of the co-translational folding (coTF) of many proteins are highly stable folding intermediates that are absent or only transiently populated off the ribosome, yet persist during translation well-beyond complete emergence of the domain from the ribosome exit tunnel. Intermediates are important for folding fidelity; however, their structures have remained elusive. Here, we have structurally characterised two coTF intermediates of an immunoglobulin-like domain by developing comprehensive ^19^F NMR analyses using chemical shifts, paramagnetic relaxation enhancement (PRE), and protein engineering. We integrated these experimental data with extensive molecular dynamics (MD) simulations to obtain atomistic structures of the folding intermediates on the ribosome. The resulting intermediate structures are distinguished by native-like folds initiated from either their N-or C-termini, and reveal parallel folding pathways, which are structurally conserved within the protein domain family, in contrast to their *in vitro* refolding mechanisms. By redirecting proteins to fold along hierarchical, parallel routes, the ribosome may promote efficient folding by avoiding kinetic traps, and regulate nascent chain assembly and targeting by auxiliary factors to maintain cellular proteostasis.

## Main

During biosynthesis, nascent proteins emerge from the ribosome and can begin to acquire their native biologically active structures^1^. The ribosome itself is thought to regulate coTF by modulating the nascent polypeptide chain’s thermodynamic stability^2–8^ and (un)folding rates^2,6,9^, through vectorial emergence^10,11^, steric restriction of the available conformational space^12^, interactions with its highly charged surface^5,13–15^ and with specific ribosomal proteins^16–18^, as well as the recruitment of molecular chaperones^19–22^. In particular, the physical chemistry of the ribosome results in the structural expansion and entropic destabilisation of unfolded nascent proteins^3^, and enthalpic destabilisation of negatively charged folded states by long-range repulsive electrostatic forces^23^. Consequently, for negatively charged proteins (∼50% proteome^24^), highly stable folding intermediates are the predominant conformer outside the ribosome exit tunnel^2,3,23^, where most tertiary folding occurs. The cellular process of coTF therefore contrasts with studies measuring the refolding of full-length denatured protein, where intermediates are unstable, and is reflected by their folding outcomes, with higher propensities to misfold or aggregate in the absence of the ribosome. Detailed descriptions of coTF pathways are absent, although structures of folding intermediates would provide experimentally accessible snapshots of *de novo* protein folding mechanisms. Such structures could therefore permit insights into how the ribosome biases nascent chain conformations away from hazardous states^25–27^ and towards conformations that are biologically active^3^ or accessible for co-translational assembly^28,29^ or chaperoning processes^19^.

Co-translational folding intermediates have generally been reported as compacted^30–32^, with some observations of native-like structure formation^2,3^ within ensemble conformations^15,33,34^. However, nascent proteins inherently interconvert between multiple conformations^2,35^, presenting a technical challenge to detailed, experimental measurements of specific intermediate state conformations. Solution-state NMR spectroscopy of ribosome-nascent chain complexes (RNCs) has enabled residue-specific level measurements of different nascent protein conformations by employing specific isotopic labelling schemes^3,5,13,16,36–38^, providing experimental data to guide the determination of accurate structures of unfolded^3,13^ and folded^23^ RNCs from all-atom MD simulations. The direct observation and resolution of multiple partially folded coTF intermediates has uniquely been enabled by ^19^F NMR^2,3,16^, which has the potential to extract structural information^39^.

Here, we have developed an array of generalisable ^19^F NMR analyses for structural biology, and obtain direct structural contacts and intramolecular distances within two distinct coTF intermediate conformations populated by an immunoglobulin-like domain on the ribosome. These ^19^F NMR experiments validate atomistic structures of the coTF intermediates, obtained using atomistic folding simulations and extensive unbiased MD simulations, and show their conservation across immunoglobulin folds. By mapping detailed coTF pathways, we thus present the structural basis of how ribosomes guide native structure acquisition of one of the commonest protein folds in the proteome.

CoTF folding intermediates identified by ^19^F NMR We previously mapped the folding free energy landscapes of the fifth filamin domain of the *Dictyostelium discoideum* gelation factor, FLN5, both off and on the ribosome, using NMR spectroscopy^2,3,5,11,36^. FLN5 is an immunoglobulin-like domain with two sandwiched anti-parallel β-sheets (**Figure 1a**). In isolation, it is a two-state folder under denaturing conditions^2^, and only weakly populates intermediates only when truncated at its C-terminus^2,11^. However, when translation-stalled on the ribosome at specific nascent chain lengths (**Figure 1b**) and fully emerged from the tunnel^36^, FLN5 populates two highly stable coTF intermediates (I1 and I2, **Figure 1d-e**) in slow exchange with the unfolded and natively folded states (exchange rate k_ex_ < 2.4 s^-1^, ref.^2^), and detected as broad NMR resonances by site-specific ^19^F-labelling using 4-trifluoromethyl-L-phenylalanine^2^ (tfmF, **Figure 1c**), and invisible by other isotopic labelling schemes^11^. Measurement of ^19^F NMR signal integrals showed that the coTF intermediates are up to 5 kcal mol^-1^ more stable than those found off the ribosome (relative to their respective unfolded states), and persist at long nascent chain lengths during translation^2^ (**Figure 1d**), despite complete availability of the domain for native folding to occur^36^. The coTF intermediates adopt a native-like hydrophobic core, evidenced by their complete destabilisation upon mutation of the natively buried residue Tyr719 (ref.^2^); however, further details of their structures are sparse.

**Figure 1.**
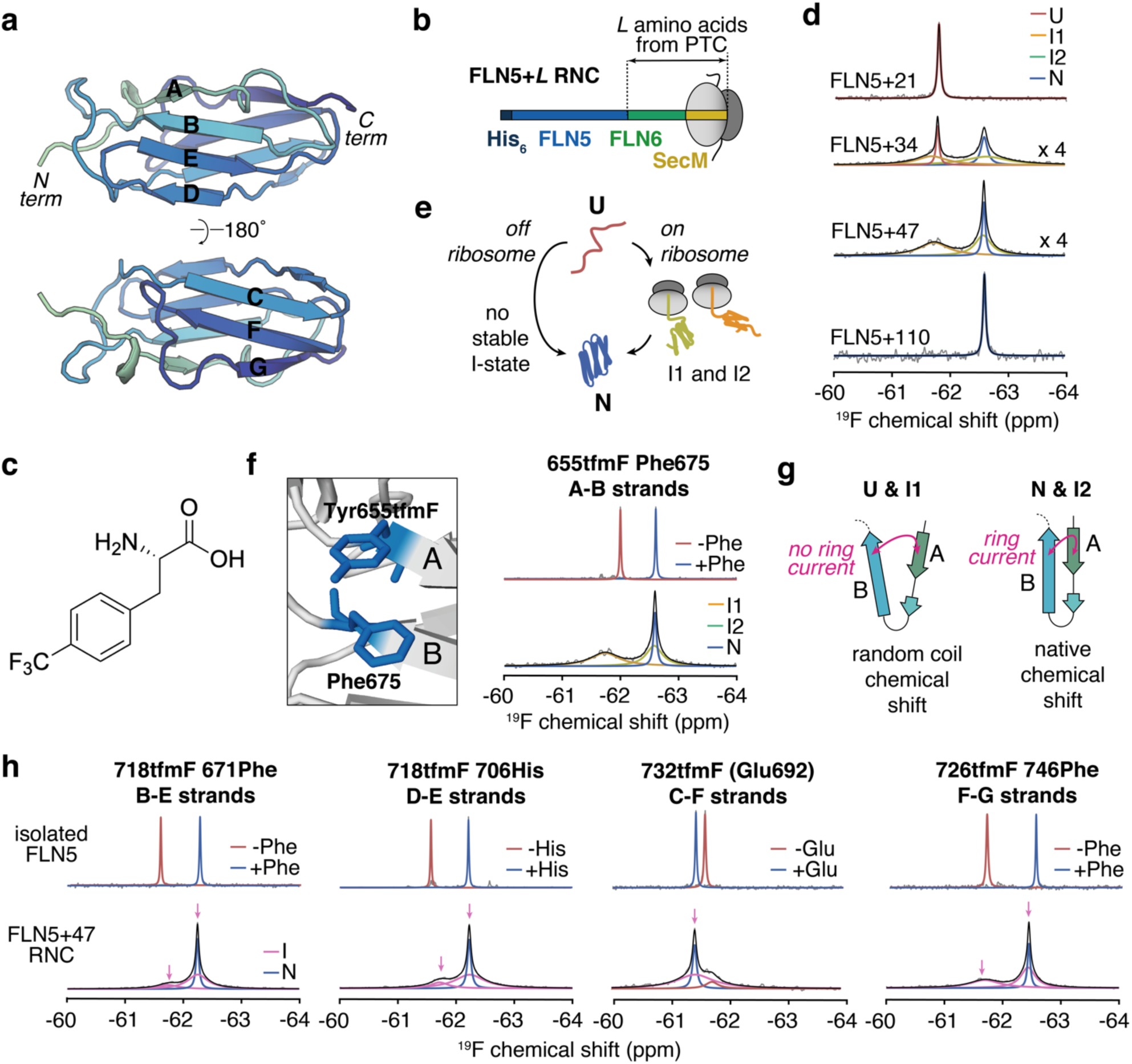
Structural characterisation of FLN5 coTF intermediates using ^19^F NMR chemical shifts. **a**, Crystal structure of FLN5 (Protein Data Bank (PDB) code 1QFH) with β-strands labelled. **b**, Schematic of FLN5 RNCs of varying *L* lengths. **c**, Chemical structure of tfmF. **d**, 1D ^19^F NMR spectra of different lengths of 655tfmF-labelled FLN5 RNCs. Observed and total fitted spectra are shown in grey and black, respectively. Native (N), unfolded (U) and intermediate state (I1, I2) resonances are indicated. Magnification of spectra by a scaling factor is indicated. **e**, Folding reactions of FLN5 on and off the ribosome. **f**, Left shows magnified view of Tyr655 and Phe675 in the FLN5 crystal structure. Right shows ^19^F NMR spectrum of 655tfmF-labelled *(top)* isolated wild-type and Phe675Ala mutant FLN5, and *(bottom)* FLN5+47 RNC. **g**, Schematic depicting ring current between two residues on adjacent β-strands and their corresponding ^19^F chemical shifts. **h**, Top shows ^19^F NMR spectra of isolated FLN5 with ^19^F-label pair, overlaid with that of isolated FLN5 tfmF-labelled without accompanying aromatic/charged residue (left to right, FLN5 718tfmF, FLN5 718tfmF, 732tfmF 692Ala, 726tfmF). Bottom shows ^19^F NMR spectra of FLN5+47 RNC with different ^19^F-label pairs. Arrows indicate chemical shifts of intermediate states. Observed and total fitted spectra are shown in grey and black, respectively. Assignment of resonances are coloured accordingly. All spectra were recorded at 500 MHz and 298 K.

Structural contacts revealed by ^19^F NMR chemical shifts To begin to extract structural information by ^19^F NMR spectroscopy, we examined ^19^F chemical shifts and their relationship to protein structure, exploiting our recent development of solvent-exposed tfmF-labels experiencing ring current effects^39^, which are distance-and angle-dependent on nearby aromatic residues. Residue 655 on the A-strand directly contacts the aromatic ring of Phe675 on the neighbouring B-strand; its tfmF chemical shift is consequently shielded from the random coil value by ∼0.8 ppm (**Figure 1f**, ref.^2^) and correlated to the ring current effect as shown by the Phe675Ala mutation^39^ (**Figure 1f**). The tfmF655 chemical shift is thus a direct reporter of the inter-residue 655-675 sidechain Van der Waals contact^39^ comprising a precise, short-range (4-6 Å) distance and angular preference, as quantifiable by structural modelling^39^. The observed broad I2 resonance of tfmF655-labelled FLN5 RNCs shares its chemical shift with the native state, and thus demonstrates a native inter-residue 655-675 tertiary contact and complete A-B strand pairing within this intermediate structure. Conversely, the random coil tfmF-655 chemical shift of I1 (-61.8 ppm) indicates the absence of the native A-B strand contact (**Figure 1f-g**).

Additional native ring currents are absent in FLN5, and tfmF-labels at other solvent-exposed sites on FLN5 yield chemical shifts that are poorly resolved between the folded and unfolded (random coil) states^39^, and thus do not show clear correlations with (local) protein structure, limiting their use in structural studies. Using ColabFold^40^, AlphaFold3 (ref.^41^) and all-atom MD simulations, we therefore rationally designed^39^ five additional ^19^F-label pairs to engineer ring currents and probe specific sidechain contacts between pairs of β-strands, encompassing the B-E, D-E, and F-G strands (**Figures 1h, S1**), akin to the introduction of (bulkier) Förster resonance energy transfer (FRET) or double electron-electron resonance (DEER) probes. Efforts to generate label pairs for C-F and A’-G using the ring current strategy resulted in more modest chemical shift dispersions insufficient to resolve broad RNC resonances (**Figure S1b**). The successful FLN5 label variants, in isolation of the ribosome, remained two-state folders and showed only mild changes to their folding free energies (ΔΔG_N-U_ < 0.8 kcal mol^-1^ relative to non-fluorinated, **Figure S1c**), being solvent-exposed; their (de)stabilising effect is expected to be largely suppressed on the RNC due to the ribosome’s mutation buffering effect^3,25^. Their ^19^F chemical shifts were substantially (de)shielded (> 0.2 ppm) from the random coil and highly sensitive to the presence of the introduced aromatic ring (**Figure 1h**), as designed. Chemical shifts engineered using ring currents can therefore be interpreted as a residue pair-specific Van der Waals contact^39^.

We introduced each label into FLN5+47 RNC, the length at which the coTF intermediates are maximally populated^2^ (44 ± 3% and 38 ± 6% of I1 and I2 respectively for FLN5+47 655tfmF, **Figure 1d**). First, FLN5+47 RNC was labelled to probe the B-E strands (718tfmF 671Phe), yielding ^19^F NMR spectra with a sharp (∼18 Hz linewidth) resonance at an identical chemical shift to that of the isolated protein (-62.2 ppm), corresponding to the native state as expected (**Figure 1d**), and two additional broad resonances at the native and random coil (-61.8 ppm) chemical shifts attributable to intermediate states (**Supplementary Note 1**). The distinct chemical shifts thus indicate two populations of intermediates, one possessing and one lacking the native B-E strand contact. Similarly, labelling of FLN5+47 RNC at an alternative position across the B-E strands (673tfmF 716His) also produced two broad resonances, having native and random coil chemical shifts (**Figure S1a**), and suggesting that the presence or absence of native contacts persists across the inter-strand network, respectively, and thus complete secondary structure formation.

When labelled to probe either the D-E (718tfmF 706His) or F-G strands (726tfmF 746Phe), FLN5 RNCs also produced broad resonances attributable to two intermediate states (**Figure 1h**). In each case, the broad resonances were found at both native and random coil chemical shifts, consistent with alternative label pairs positioned along the same strands (710tfmF 716His, and 728tfmF 744His, **Figure S1a**, **Supplementary Note 1**). In addition to the findings for the A-B and B-E strands, native inter-strand contacts across the D-E and F-G strands are therefore only found in one intermediate.

The 732tfmF (F-strand) chemical shift is highly sensitive to the electrostatic charge of the native neighbouring residue Glu692 (C-strand) in the isolated protein^39^ (**Figures 1h, S1a, S1e**). In addition to the native state resonance, the ^19^F NMR spectrum of FLN5+47 732tfmF showed a single broad resonance (67 ± 11% population) observed at the same chemical shift that was attributable to an intermediate state (**Figure S1a**). Only a lowly populated (11 ± 4%) peak was observed at the random coil chemical shift, having a linewidth (**Table S1**) and small population (**Table S2**) consistent with an unfolded state, the latter as expected from the more destabilising nature of the 732tfmF-label (relative to other ^19^F-label pairs, **Figure S1c**). Similar results were found for the shorter FLN5+34 RNC (**Figure S1a**, **Supplementary Note 1**). These data indicate that both I1 and I2 intermediates possess a native 732-692 contact. Additional structural contacts in the hydrophobic core were measured with natively buried ^19^F-labels (**Figure S2**).

Collectively, by using and engineering ^19^F chemical shifts that unambiguously probe specific intramolecular contacts, we resolved two distinct intermediate states in slow exchange with the natively folded state (**Supplementary Note 1**), both with native-like structure between the C-F strands, and where additional native contacts across the remaining β-strand pairs and buried core are found in only one intermediate conformation.

Engineering reversible cross-links stabilises I1

Equipped with NMR data probing structural features across the intermediate states, we next aimed to attribute the broad resonances to either I1 or I2 to generate a dataset of chemical shifts (i.e. structural contacts) specific to either coTF intermediate. To achieve this, we sought to selectively (de)stabilise one intermediate conformation. Using principles of psi-value analysis^42^, we introduced reversible cross-links by engineering bi-histidine metal ion binding sites to stabilise specific β-strands. We experimentally screened for potential inter-strand binding sites by assessing different bi-histidine mutants of isolated FLN5 and their stabilities in increasing concentrations of a range of different metals (**Figure S3a**). Of the seven mutants tested, one variant, 726His 746His (F-G strands), showed a progressive stabilisation of up to-1.45 kcal mol^-1^ in its folding free energy with increasing concentrations of NiCl_2_ (**Figures 2a-c, S3c**). The metal binding site was confirmed by ^1^H,^15^N-NMR, where addition of the paramagnetic Ni(II) metal resulted in increased line broadening localised at the mutation sites (**Figures 2a, S3b-h**), in accord with AlphaFold3 predictions showing coordination between the metal ion and the protein via the engineered histidines and the neighbouring residue Asp744 (**Figure 2a**).

**Figure 2.**
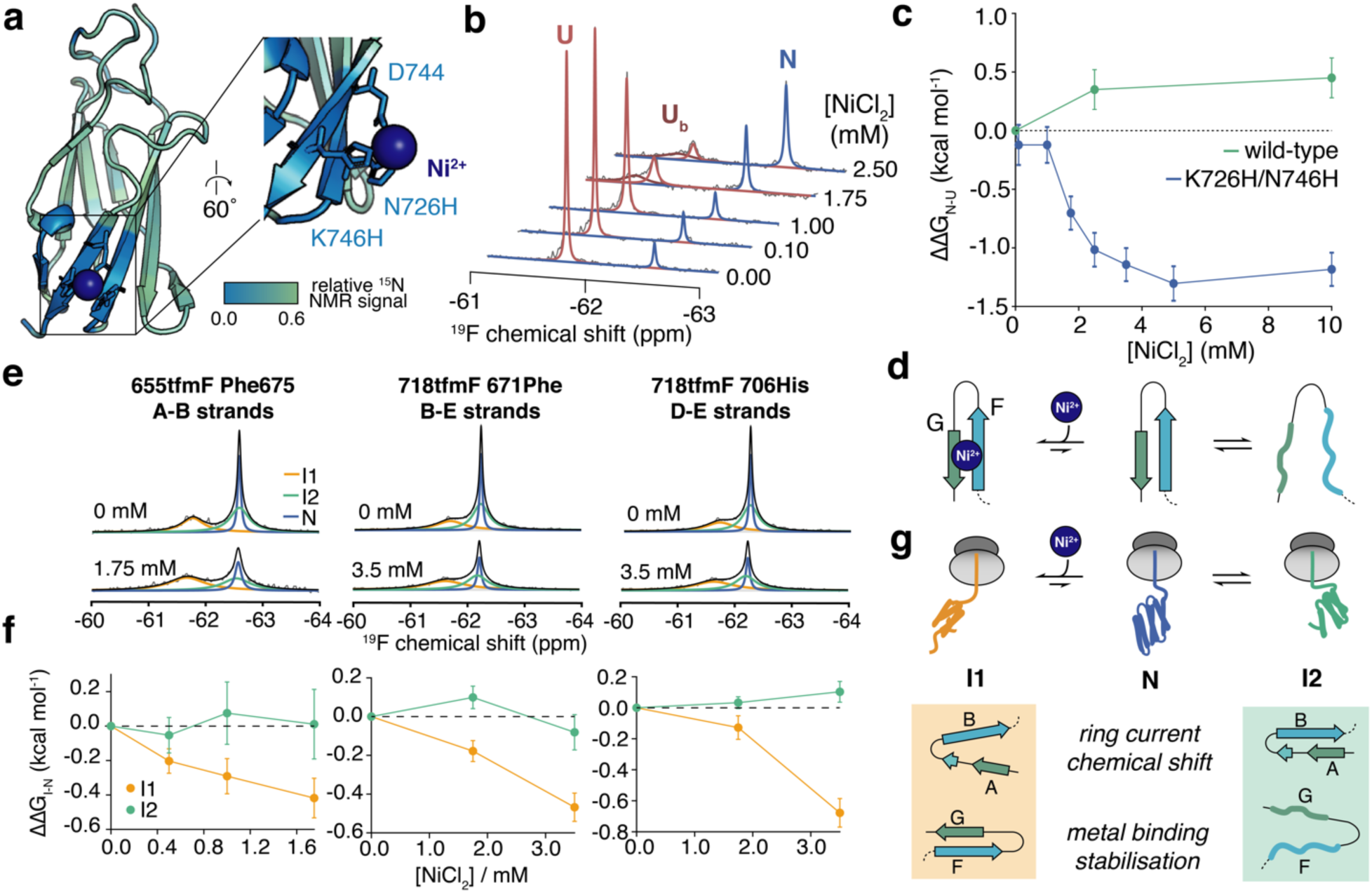
Stabilising a specific coTF intermediate using an engineered ligand binding site. **a**, AlphaFold3 predicted structure of FLN5 Asn726His Lys746His in complex with a metal ion, coloured according to the ^1^H,^15^N-correlated NMR signal intensity in 2.5 mM NiCl_2_ relative to without NiCl_2_ (**Figure S3c** for complete dataset). **b**, ^19^F NMR spectra of 655tfmF-labelled FLN5 Asn726His Lys746His incubated in 2 M guanidine hydrochloride and varying NiCl_2_ concentrations. Observed spectra are shown in grey. Unfolded (U), nickel-bound unfolded (U_b_, see **Figure S3g**), and native (N) states are in colour and indicated. **c**, Folding free energy changes of isolated wild-type and bi-histidine FLN5 in varying NiCl_2_ concentrations (relative to without NiCl_2_), determined by lineshape analysis of spectra shown in **b** and **Figure S3c**. Errors were determined by bootstrapping of residuals from lineshape fits**. d**, Schematic of equilibrium reaction between unfolded and folded β-strand pairs titrated with Ni(II). **e**, ^19^F NMR spectra of FLN5+47 Asn726His Lys746His RNCs tfmF-labelled at different positions, with and without 3.5 mM NiCl_2_ (**Figure S3j** for complete titration data). Observed and total fitted spectra are shown in grey and black, respectively. Assignment of resonances are coloured accordingly. **f**, Changes in Gibbs free energies of intermediate relative to native state, determined by lineshape analysis of spectra shown in e and Figure S3j (**Figure S3j** for complete titration data). Errors were determined by bootstrapping of residuals from lineshape fits**. g**, Schematic of equilibrium reaction between I1, I2 and N titrated with Ni(II). Nickel-binding stabilises I1 more than I2, inferring the presence and absence of the bi-histidine binding site, respectively. I1 and I2 are also distinguished by their ring current-sensitive chemical shifts. All spectra were recorded at 500 MHz and 298 K.

Having designed a conformation-specific, concentration-dependent ligand binding site (**Figure 2d**), we next introduced the bi-histidine mutations into FLN5+47 RNCs (**Figures 1h, S3i-k**). FLN5+47 tfmF655 RNCs (probing A-B strands, **Figure 1f**) yielded resonances corresponding to I1, I2, and N as expected (**Figure 2e**). Addition of increasing amounts of Ni(II) to the RNC sample resulted in increasingly broader resonances, as anticipated from the paramagnetic effect (**Figure S3k**), but critically also a progressive shift in the folding equilibrium towards the I1 state (**Figures 2e-f**, **S3j**), only in the presence of the bi-histidine mutations (**Figure S3j**). An analysis of the signal integrals to determine the populations and Gibb’s free energies of each state revealed that I1 was stabilised more than I2 (each relative to N, **Figure 2f**). Given the distinct chemical shifts obtained for the F-G strand ^19^F-labels (**Figure 1h**, **S1a**), this indicated the presence of a folded binding site across the F-G strands in I1 that was likely absent in I2 (**Figure 2g**). I1 therefore possesses a native F-G strand conformation, but lacks A-B strand contacts (**Figure 2g**), while I2 exhibits the opposite.

When the bi-histidine site was introduced in FLN5+47 RNCs possessing the remaining ^19^F-label pairs that showed two distinct intermediate resonances (**Figures 1h, S1a**), one intermediate (I1) was consistently stabilised more than the other (I2) when incubated with Ni(II) (**Figures 2e-f, S3j**). These data permitted us to assign the chemical shifts across all the tfmF-labels to either the I1 or I2 intermediate (**Figures 1g, S3j**). Collectively, they show I1 likely possesses a natively folded sheet 2 (CFG strands) with native contacts absent across sheet 1 (ABED strands). However, native-like strands are likely to be at least partially formed within sheet 1, since I1 is completely destabilised by the natively buried Tyr719Glu mutation in the core between the two β-sheets^2^. Meanwhile, on the basis of the ^19^F NMR contacts, I2 appears to closely resemble the intermediate found off in isolated C-terminally truncated variants of FLN5 (ref.^11^), comprising a natively folded A-F strand structure with a disordered C-terminal G-strand; accordingly, native-like chemical shifts for the natively buried sites tfmF715 and tfmF727 also correspond to the I2 intermediate (attributed using the same bi-histidine binding approach, **Figures S2a, S3j-k**).

Structures of folding intermediates on the ribosome Next, we aimed to produce atomistic structural ensembles of the FLN5+47 RNC intermediate states. The structural similarity of I2 to the isolated, truncated intermediate enabled us to produce a starting model of the RNC, by attaching the previously determined structure of the latter^11^ to the ribosome (**Supplementary Note 2**). However, in the absence of any analogous structures to I1, we sought to simulate coTF and produce detailed pathways from the unfolded to natively folded state using an all-atom (including ribosome, ions, and solvent) ratchet-and-pawl MD (rMD) approach (**Supplementary Note 2**, **Supplementary Videos 1-2**, **Figures S4-6**). The rMD simulations enabled us to successfully capture folding events, which experimentally occur on the timescale of > 0.8 s^-1^ (ref.^2^), beyond the reach of conventional MD simulations. From the obtained coTF trajectories, we identified one on-pathway folding intermediate state (**Figure S5**), which from our experimental dataset appeared to possess structural features common to that of I1. To assess this further, we subsequently subjected the putative I1 and I2 models to extensive atomistic unbiased MD simulations. We found their folded regions to be stable on the microsecond timescale (**Figure 3**, **Supplementary Videos 3-4**), and therefore compared the atomistic structural ensembles against those previously obtained for the natively folded RNC^23^.

**Figure 3.**
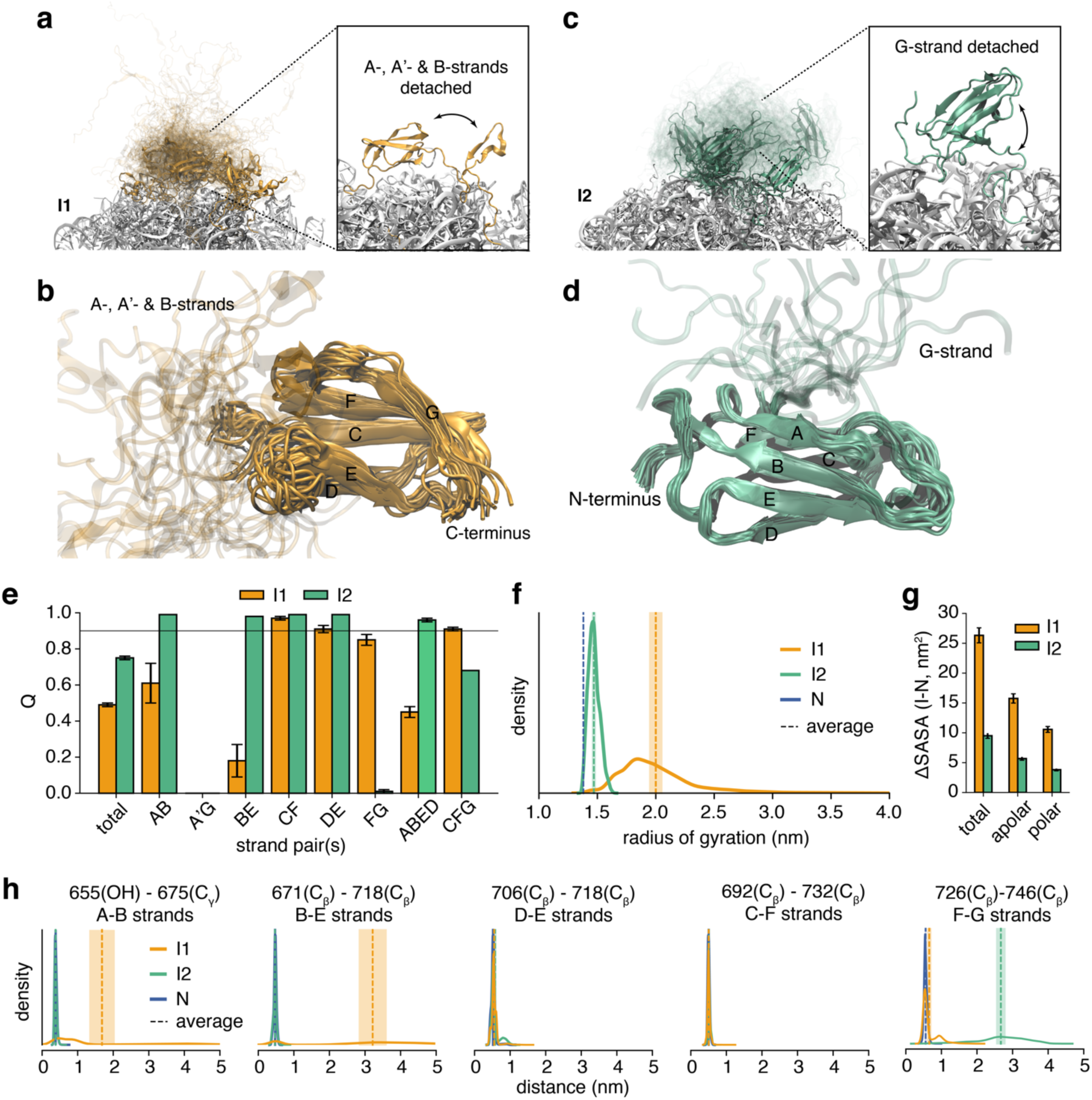
Atomistic structures of FLN5+47 coTF intermediates on the ribosome. **a**, Structural ensemble of FLN5+47 I1 on the ribosome, inset shows a representative structure. **b**, Lowest energy structural cluster of I1 (**Table S4**) aligned against the common C-G strand core. **c**, Structural ensemble of FLN5+47 I2 on the ribosome, inset shows a representative structure. **b**, Lowest energy structural cluster of I2 (**Table S4**) aligned against the common A-F strand core. **e**, Average fraction of native contacts (Q) of FLN5 and across different β-strand pairs and β-sheets for I1 and I2 (mean ± s.e.m. from 15 independent simulations of 1 µs). **f**, Radius of gyration probability distribution and mean ± s.e.m. of I1, I2 and native states. **g**, Difference in solvent accessible surface area (SASA) of I1 and I2 relative to the native state. **h**, Distance distributions and mean average ± s.e.m. of residue sidechains corresponding to ^19^F-label pairs calculated for I1, I2 and native state structural ensembles.

The structures of the putative I1 state (**Figures 3a-b**, **S6a-g**) possess a native-like C-G strand folded core (**Figure 3e-f**); the remaining N-terminal A-, A’-and B-strands are largely detached from the structure, and form either a completely disordered tail, or a β-hairpin (**Figure 3e, S6b**). The loss of the A-B strands from the folded structure results in transient non-native contacts, predominantly between the β-sheets at the exposed E-G and A’-F strand interfaces (∼20-30%, **Figure S6c-e**). However, we also find a subpopulation (∼24%) where the B-strand folds onto the C-G core, with the A-A’ strands remaining disordered (**Figure S6b**); this conformational heterogeneity shows multiple accessible pathways from I1 to the native state, as also reflected in our folding simulations (**Figure S5**). Overall, detachment of the N-terminal strands causes an expansion of the structural ensemble, with a larger radius of gyration and increased solvent accessible surface area (SASA) compared to the native state (**Figure 3g**).

Structures of the putative I2 state (**Figures 3c-d, S6a-g**) adopt a more ordered and compact native-like conformation, with a smaller increase in SASA (**Figures 3f-g**). Native contacts are found across the A-F strands (**Figure 3e**), with a non-native trans-Pro742 and disordered G-strand (residues 743-747) detached from the rest of the domain permitting some non-native interactions between A’-F strands (**Figure S6c-f**). We note that energetic calculations show native G-strand folding is dependent on the cis-trans isomerisation state of Pro742 (**Supplementary Note 2, Figure 4**).

**Figure 4.**
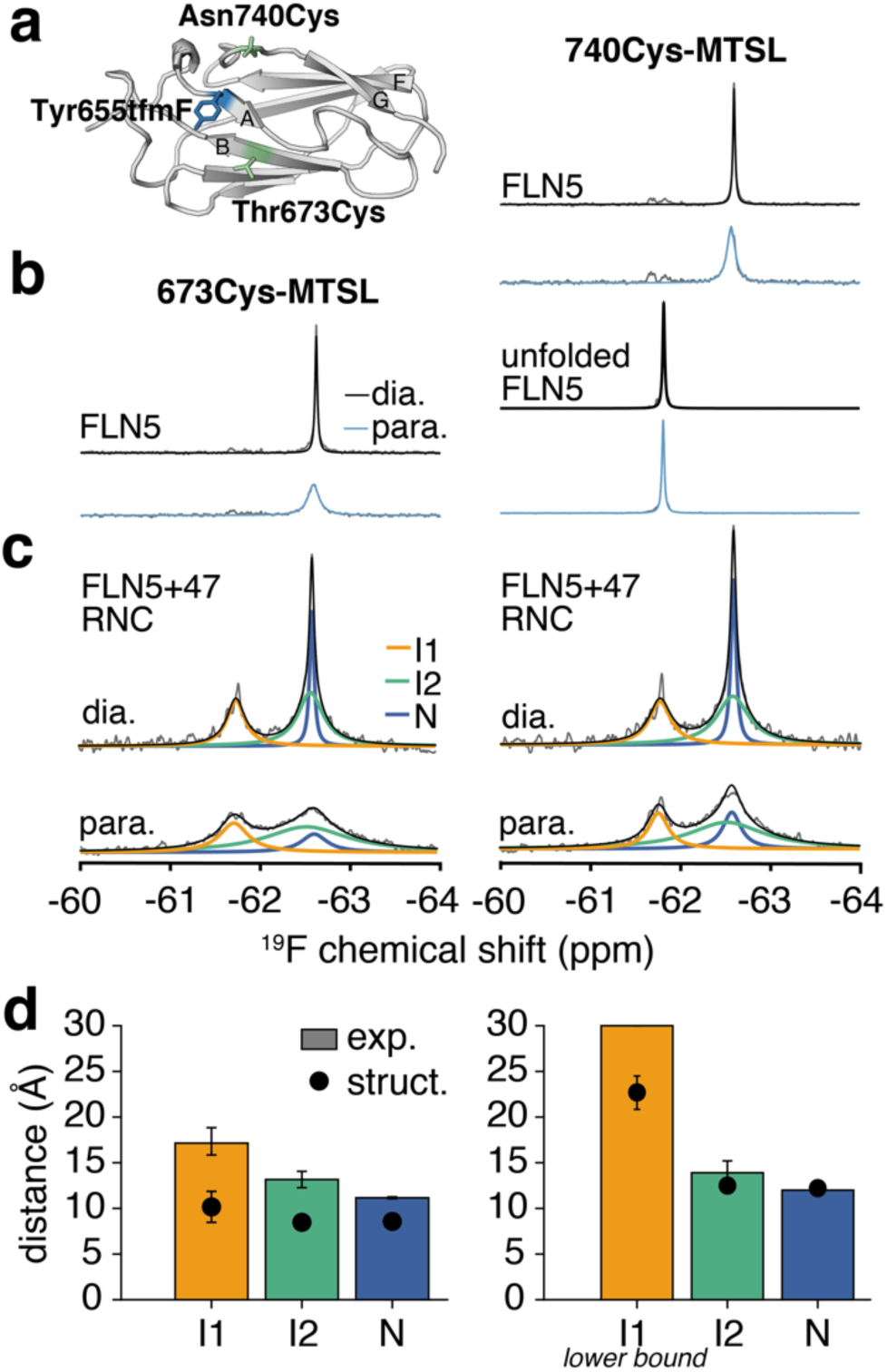
Validation of coTF intermediate structures by ^19^F PRE NMR. **a**, Location of tfmF-and paramagnetic spin-labels shown on the FLN5 crystal structure (PDB: 1QFH). **b**, ^19^F NMR spectra of isolated 655tfmF-labelled FLN5 (pseudo-wild type Cys747Val (**Methods**), and unfolded with destabilising mutations (A_3_A_3_, ref.^36^)), spin-labelled at position *(left)* 673Cys and *(right)* 740Cys. Observed spectra are shown in grey. **c**, ^19^F NMR spectra of 655tfmF-labelled FLN5+47 RNC (pseudo-wild type Cys747Val), spin-labelled at position *(left)* 673Cys and *(right)* 740Cys. Observed and total fitted spectra are shown in grey and black, respectively. Assignment of resonances are coloured accordingly**. d**, Distances between the ^19^F-and spin-labels, bars show experimental values determined by lineshape analysis of spectra shown in **c** (**Methods**), dots show *r*^6^-averaged values (mean ± s.e.m.) from the intermediate structures. Errors of experimental values determined by bootstrapping of residuals from lineshape fits. All spectra were recorded at 500 MHz and 298 K.

To assess whether the obtained structures faithfully represent our experimental data (**Figures 1-2**), we examined their intramolecular contacts and compared them against the obtained ^19^F chemical shifts, since we engineered the latter to be sensitive reporters of precise, short-range inter-residue distances^39^. We calculated distance distributions between the pairs of residues labelled for NMR analysis (**Methods**, **Figure 3h, S6j**), and found that all could account for the observed chemical shifts of I2: those with a native chemical shift showed short (∼0.5 nm), native-like inter-residue distances, consistent with the measured van der Waal’s contact, while residues labelling the F-G strands were distanced far (>2 nm) from each other, as reflected by their random coil chemical shifts (**Figures 1f, 1h, S1a**). Analyses for I1 also showed good agreement between inter-residue distances and their ring-current induced chemical shifts (**Figures 3d, S6j**). The exceptions were the two D-E strand labels, but which could instead be resolved and rationalised using more precise, explicit modelling of the ^19^F-label pair: the D-E strand pairs form native backbone contacts but lack sidechain interactions in line with the random coil chemical shifts found in this region (**Supplementary Note 3**, **Figure S3l**). Likewise, such simulations entirely validate residue-contacts in the core of the domain (**Supplementary Note 3**, **Figure S6m-n**). In summary, our residue-contact analyses show that both I1 and I2 structures account for all of our ^19^F chemical shift data, and thus also that the rMD folding simulations captured an intermediate conformation corresponding to I1.

Interactions and orientations of intermediates with the ribosome The intermediate state structures show transient interactions with the ribosome surface, predominantly with ribosomal proteins uL24, uL29 and RNA H59 lining the vestibule of the exit tunnel, and more favourably than the native state (**Figure S6h-i**). Their consequently slower calculated rotational correlation times (**Figure S6k**) are reflected experimentally by the increased line broadening of their ^19^F NMR resonances^2,3^ (relative to the native state, **Table S1**). Locally enhanced binding occurs at their more positively charged N-termini and the loop between the B-C strands, biasing their orientations with the ribosome (**Figure S6h**). This observation is consistent with previous experimental measurements of FLN5 RNCs bearing charge reversal mutations within the two binding sites (Lys646Glu Lys680Glu) showing thermodynamic destabilisation of both I1 and I2 (ref.^2^). The detached C-terminal G-strand in I2 also interacts more strongly with the ribosome surface (**Figure S6h**), a region that has been experimentally shown to bind and stabilise unfolded FLN5 on the ribosome^5^. We explored the presence of intermolecular interactions further by directly examining specific contacts between nascent chain residues and ribosomal protein uL24 that lines the exit tunnel using biochemical cross-linking assays. These experiments showed good agreement between (un)observed crosslinks, and intermolecular contacts calculated from the structural ensembles (**Supplementary Note 5**, **Figure S8d-e**).

Our coTF intermediate structural ensembles were further assessed against our previously obtained cryogenic electron microscopy (cryo-EM) maps of FLN5+47 RNC (Mitropoulou *et al.*, in preparation). We performed two orthogonal analyses: firstly by determining cross-correlations of single structures within our intermediate ensembles to the cryo-EM densities, and secondly by producing structural ensembles collectively combining I1, I2 and native state structures (this work, and ref.^23^ respectively) reweighted by the cryo-EM densities^43^ (**Supplementary Note 5, Figures S8a-c)**. Both approaches showed quantitative improvements to the native state fits when incorporating the intermediate states, and we thus conclude that the cryo-EM maps support the determined intermediate models.

^19^F PRE NMR measurements validate intermediate structures We sought to validate our determined intermediate structures and complement the ^19^F chemical shift data by measuring long-range (10-30 Å) distances using ^19^F PRE NMR. Having identified detachment of the A-B strands as a key structural feature in I1 (**Figure 3a-b**), we covalently linked paramagnetic spin labels on the adjacent B-strand (Thr673Cys) and F-G loop (Asn740Cys), and measured their effect on the tfmF655 (A-strand) resonance (**Methods**, **Figure 4a**). As expected, the isolated, folded protein showed line broadening in the presence of either paramagnetic label, but only marginal changes when the protein was fully unfolded (**Figures 4b, S1a**).

The native state resonance was also broadened in paramagnetic spin-labelled FLN5+47 RNC. However, lineshape analysis of the resonances (**Methods**) revealed differential effects between the I1 and I2 linewidths and the two spin label sites (**Figures 4c**, **S7b-d**). We used the measured linewidths to calculate distances between the ^19^F-and paramagnetic spin labels (**Supplementary Note 4**). Native-like distances between A-B strands and A-loop(FG) were evaluated for I2 (**Figure 4d**), consistent with its determined structural ensemble (**Figure 3c-d**). In contrast, larger distances were calculated for I1: in particular, a substantial increase (to > 20 Å) for the 740Cys-MTSL spin label (measuring between A-B strands and A-loop(FG)) compared to the native state (**Figure 4d**). These data agree with the structures (**Figure 3a-b**), where the A-strand remains detached from the C-terminus across all conformations, while the formation of a β-hairpin in subpopulations of I1 rationalises the shorter ensemble-averaged distance between A-B strands.

Thus collectively, the NMR chemical shifts (**Figures 1-2**), cross-linking (**Figure S8d**), PRE (**Figures 4**, **S7**) and cryo-EM data (**Figure 8a-c**) confirm that the determined structures of I1 and I2 are highly representative of the experimental data.

Conservation of intermediate structures across filamin domains Finally, we sought to examine whether the intermediates of FLN5 are conserved across the coTF pathways of domains with a similar fold. We therefore studied the folding of three additional immunoglobulin-like domains (of a multi-domain protein) on the ribosome (**Figure 5a**): FLN4, the domain preceding FLN5 (47% identity relative to FLN5); FLNa21, the 21^st^ domain of human filamin A (the human homologue of gelation factor, 31% identity); and I27, the 27^th^ immunoglobulin-like domain of titin (24% identity), whose smaller size (89 vs 105 residues compared to FLN5) is accompanied by a subtly different topology. For each protein, we rationally designed and experimentally validated ^19^F-label pairs with ring currents in the natively folded state^39^, permitting us to assess inter-strand contacts across each immunoglobulin-like fold.

**Figure 5.**
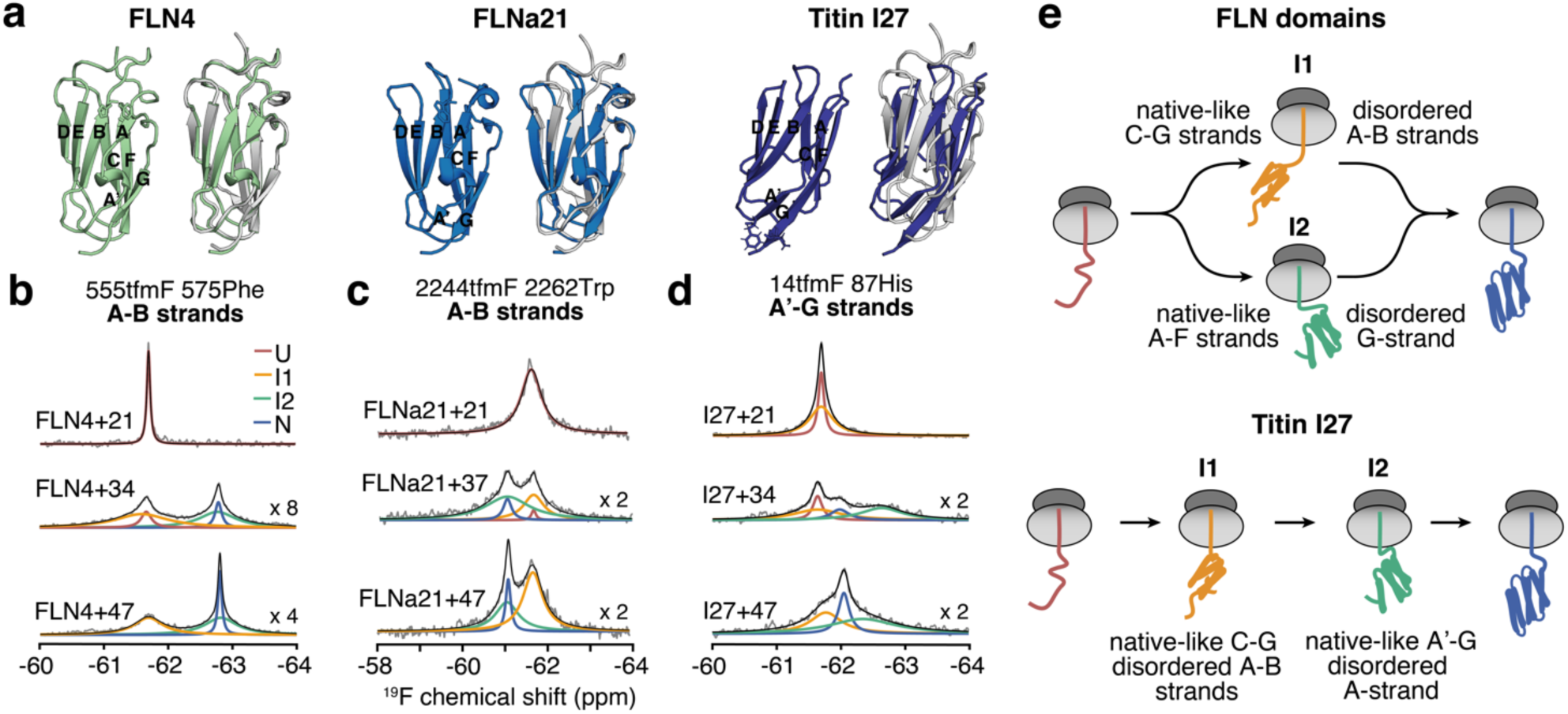
^19^F NMR of RNCs of immunoglobulin domains show conservation of filamin coTF pathways. a, Structures of FLN4 (AlphaFold3), FLNa21 (PDB: 2W0P), and titin I27 (PDB: 1TIT), and alignment with FLN5 structure (PDB: 1QFH). **b-d**, ^19^F NMR spectra of 555tfmF 575His-labelled FLN4+*L*, 2244tfmF 2262Trp-labelled FLNa21+*L*, and 14tfmF 87His-labelled I27+*L* RNCs, with linkers of *L* amino acids deriving from the subsequent domain. Observed and total fitted spectra are shown in grey and black, respectively. Assignment of resonances are coloured accordingly. Magnification of spectra by a scaling factor is indicated. **e**, Summary of ^19^F NMR data of other labelling sites, shown as schematic of coTF intermediates and pathways of the filamins (FLN4, FLN5, and FLNa21) and I27 (all NMR data shown in **Figure S9a-c**). All spectra were recorded at 500 MHz and 298 K.

We first aimed to identify nascent chain conformations populated on the ribosome, and therefore produced a library of RNCs varying in length *L* of the subsequent domain (domain+*L* RNC). FLN4 RNCs ^19^F-labelled at the A-B strands (555tfmF 575Phe) showed a sharp resonance with a random coil chemical shift at the shortest RNC length (FLN4+21), which reduced in population with increasing RNC length, and thus attributable to the unfolded state; concurrently, a sharp resonance with a native chemical shift progressively increased in signal intensity and was ascribed to the natively folded state (**Figure 5b**, **S9a**). Importantly, two broad resonances, maximally populated at intermediate RNC lengths, were also found at each of the same chemical shifts (**Figure 5b**, **S9a**). Thus, identically to FLN5 (**Figure 1f**), FLN4 populates two coTF intermediates on the ribosome, distinguished by the presence of A-B strand contacts. Indeed, analysis using other ^19^F-label pairs (B-E, F-G strands) indicate that the same inter-residue contacts are found in FLN4 as in FLN5 coTF intermediates (**Figure 5e**, **S9a**). We also probed the A’-G strand, using a uniquely successful ^19^F-label pair in FLN4 (565tfmF 646His, **Figure S9a**), and found the absence of contacts for either intermediate state; these data are consistent with the determined structures of FLN5 I1 and I2, having detached A’-and G-strands respectively (**Figure 3a-d**). Similar ^19^F NMR analyses of FLNa21 RNCs (**Figures 5c, 5e, S9b**) also conclude the population of two coTF intermediates with the same structural features as those of FLN5. We therefore find that the determined structures of FLN5 coTF intermediates are conserved across the folding of filamin domains.

We have previously identified two coTF intermediates of titin I27 on the ribosome^3^, and show here that their populations alter with RNC length, as expected (**Figures 5a, 9c**). However, in contrast to the filamin domains, ^19^F NMR spectra of I27 RNCs labelled across the A’-G strands (14tfmF 87His) reveal an intermediate state resonance (I1) at the random coil chemical shift, but interestingly also at a significantly more shielded chemical shift to that of the native state (I2). A weakly populated intermediate having the same chemical shift was also found in a destabilised variant of I27 (Phe73Ala) in isolation^3^ (**Figure S9c**), suggesting that its conformation was similar to the ribosome-bound I2 intermediate, and whose structural features have previously been examined^44,45^. Indeed, all-atom MD simulations of the isolated I27 intermediate reconcile the observed chemical shifts and contacts of the I2 intermediate (**Supplementary Note 6**). Overall, the I27 coTF intermediates I1 and I2 are defined by detached A-B and A-strands (respectively) from a common native-like β-sandwich core, comprised of β-strands C-G (**Figure 5e**). Thus, while the filamin I1 state is likely conserved, I2 is subtly distinct to those adopted by filamin domains.

## Discussion

We have developed a range of ^19^F NMR analyses and used these together with MD simulations to determine the first atomistic structural ensembles of coTF intermediates on the ribosome. A particular strength of ^19^F NMR, and crucial to obtaining direct, precise structural contacts, is its ability to resolve interconverting conformers^2^. We extracted short-and long-range distances from ^19^F NMR chemical shifts (**Figures 1, S1-2**) and PRE measurements (**Figures 4, S7-8**) respectively, and obtained systematic structural analyses specific to each intermediate state using a psi-value-based analysis (**Figures 2, S3**). Our atomistic simulations of FLN5 on the ribosome reveal stable, partially folded intermediate state conformations with common structural features (**Figure 3**), which show striking and entire consistency with the experimental NMR, cryo-EM, and biochemical data (**Figure S8**). Overall, our study advances the utility of ^19^F NMR in structural studies, and establishes a generalisable approach to determine and validate atomistic structures of large macromolecular complexes.

Our structural ensembles of I1 and I2 show a native-like, reduced β-sandwich core, with detached A-B or G strands indicative of folding initiating at either the C-or N-terminus respectively. Thus, the two coTF intermediates are likely populated along parallel, rather than sequential, folding pathways. Indeed, only one intermediate is populated along the same folding trajectory sampled by rMD simulations (**Figure S5**), while NMR data show that both intermediates are simultaneously, rather than successively, populated at varying RNC lengths^2^ (**Figure 1d**). We note that energetic calculations (**Figure S4**) and NMR measurements^11^ show G-strand folding is dependent on the isomerisation state of the preceding Pro742, which adopts the trans conformation in I2 with a disordered G-strand, but a cis conformation in I1 and N where the G-strands are natively folded (**Figure 3**); thus, the cis-proline likely acts as a molecular switch of the conformational state of the C-terminal strand^11^ (**Figure S5**).

Notably, both the natively cis-Pro742 (ref.^11^) and coTF pathways are conserved across filamin domains (**Figure 5**). In contrast, titin I27 appears to likely fold via two sequential intermediates instead; its structure differs from the filamins (**Figure 5a**), having a smaller size, different topology, and lacking the conserved cis-proline. Thus, the parallel folding pathways identified are (only) conserved across filamins. This observation is reminiscent of the ribosome’s ability to thermodynamically^3^ and structurally^25^ buffer the effects of (de)stabilising point mutations (FLN4 or FLNa21 being substantially mutated variants of FLN5). The presence of parallel pathways on the ribosome via two possible intermediates has previously been inferred^46^, and could provide redundancy of mechanistic routes towards the native state, to improve folding efficiencies and provide resistance against external perturbations. Indeed, the rMD and structural ensemble of I1 reveal two additional pathways to the native fold, either via attachment of an A-B strand β-hairpin, or successive folding of the B-and A-strands to the folded core. Furthermore, the conserved coTF folding pathways are in contrast to un/refolding experiments of the isolated proteins^47^, which have shown distinct mechanisms for FLN4 and FLN5, indicating that the ribosome guides folding of similar protein structures via a common, generalised pathway. Interestingly, I27 has been reported to refold via two distinct, parallel transition states and pathways off, in contrast to a sequential pathway on the ribosome (this work), and whose flux is altered with denaturant concentration and mutations^48^; in a similar manner, the ribosome appears to alter the flux towards specific protein folding pathways.

The long-lived coTF intermediates appear to share structural similarities to those found transiently off the ribosome by FLN4 (ref.^47,49^), N-terminal fragments of FLN5 (ref.^2,11^), I27 (ref.^44,45^), and those of refolding transition states of immunoglobulin-like domains^50^. This observation suggests that isolated intermediate conformations are explored on the ribosome, but novel conformational states not observed in isolation also become thermodynamically accessible. The thermodynamic basis for stable coTF intermediates originates from the destabilisation of both the unfolded^3^ and folded states^23^, and so hierarchical folding via partially folded intermediates is likely one means to minimise repulsive negatively charged globular structures near the ribosome surface (**Table S5**). Our structures also highlight the range of intermediate conformational states that can be stabilised. The high degree of disorder in the I1 coTF intermediate indicates that conformational entropy substantially contributes to its thermodynamic stability. In contrast, the stability of the more native-like I2 structure is likely to benefit from larger enthalpic contributions from intra-and intermolecular interactions. The distinct structures of I1 and I2, yet similar folding free energies^2^, indicate that the loss of extensive solvation of the unfolded state to form (partially) folded structures can be compensated through a complex array of different entropic or enthalpic contributions.

The thermodynamic stabilisation of such intermediates on the ribosome persists for long durations of translation^2^, and for FLN4, whose intermediates are obligate for folding^47^, even after translation of the subsequent domain (**Figure S9a**), ensuring step-wise, hierarchical acquisition of native structure under quasi-equilibrium conditions^32^. In the context of multi-domain folding, the population of structurally expanded intermediate states of one domain may be advantageous to the thermodynamic stability of the preceding (fully translated) domain, since this would effectively increase the distance of the preceding domain from the destabilising, repulsive ribosome surface^23^, while also limiting exposure of misfolding-prone unfolded states. Such prolonged lifetimes of intermediates, in contrast to highly transient populations off the ribosome, may also provide an opportunity for chaperones^19–21^, co-factors^51^, enzymes, and quality control machinery^52^ to survey, engage, and modify^53^ specific nascent chain conformations, or for recruitment of other nascent proteins during co-translational assembly^28,29^. Intriguingly, filamin function is highly regulated by serine phosphorylation at consensus sequences in humans^54^, comprising conserved, natively-inaccessible serine-cis-proline motifs preceding the G-strand. Strand-detachment and a trans-proline in the I2 intermediate exposes the phosphorylation site, potentially implicating the formation of stable coTF intermediates to co-translational modifications^55^, and could therefore alter the domain’s post-translational function by stabilising and trapping the domain in its phosphorylated I2 state.

Future detailed characterisations of nascent chain structure will permit mechanistic insights into folding pathways of other protein folds, misfolding^25,27,56,57^ and other co-translational events, offering prospects for both deeper structural insights and targeted modulation of such processes with therapeutics.

## Methods

### Sample preparation

DNA constructs of FLN5, FLN4, FLNa21, and titin I27 were previously described^2,3,36,39^. Mutations and amber stop codons were introduced using standard site-directed mutagenesis procedures, and constructs verified by DNA sequencing. For PRE NMR experiments, the additional mutation Cys747Val was introduced to yield a cysteine-free background for site-specific spin-labelling, as previously described^3^. Isolated proteins were expressed as His-tagged proteins (FLNa21 as GST-fused protein^39^), isotopically labelled in *Escherichia coli* BL21 DE3-Gold cells (using amber suppression for incorporation of tfmF), and purified as previously described^2,36,39^. RNC constructs comprised a His-tag for purification and an arrest-enhanced variant of the SecM stalling sequence, FSTPVWIWWWPRIRGPP, as previously described^2^. RNCs were expressed, isotopically labelled, and purified as previously described^2^. For inter-molecular cross-linking experiments, RNCs were expressed in CRISPR-modified *E. coli* BL21 strains with cysteine mutations in uL24, as previously described^3^. MTSL spin-labels for PRE NMR were introduced into purified samples, as previously described^3^. Sample purity and integrity were assessed using SDS-PAGE and western blot analyses with an anti-hexahistidine horseradish peroxide-conjugated antibody (Invitrogen, 1:5000 dilution), as previously described^58^.

### NMR spectroscopy

NMR samples were prepared in Tico buffer (10 mM Hepes, 30 mM NH_4_Cl, 12 mM MgCl_2_, 1 mM EDTA), supplemented with 10% (v/v) D_2_O and 0.001% (w/v) DSS as a reference. NMR data were recorded using a 500-(^19^F and ^1^H,^15^N NMR) or 800-MHz (^1^H,^15^N NMR) Bruker Avance III spectrometer, both equipped with a TCI cryoprobe, and Topspin 3.5pl2, and at 298 K, unless stated otherwise. Data were processed using NMRPipe (v11.7)^59^, CCPN (v2.4), MATLAB (R2017a, Mathworks), and Julia 1.5 (ref.^60^).

1D ^19^F pulse-acquire experiments were recorded using a 350-ms acquisition time and 1.5-s recycle delay. Multiple short experiments were recorded in succession to monitor changes in RNC sample integrity, and data indicating nascent chain release or sample degradation through changes in linewidths, signal intensities or chemical shifts were not used to produce the final summed spectrum, as previously described^2,3^. Processed spectra were baseline-corrected, and peaks fit to Lorentzian lineshapes, with errors of the linewidths and integrals (populations) estimated using bootstrapping (200 iterations, calculating the standard error of the mean), as previously described^2^. The time-domain ^19^F NMR spectra were multiplied with an exponential window function with a line broadening factor of 5 (isolated protein) or 10 Hz (RNC), unless stated otherwise, prior to Fourier transformation. Integrals were used to calculate folding free energies using Δ*G* = -*RT*ln(*K*_eq_). 2D ^1^H,^15^N SOFAST-HMQC experiments^61^ were recorded using an acquisition time of 50 ms in the direct dimension, and inter-scan delays of 100 ms. Cosine-squared window functions were used to process the spectra.

^19^F PRE rates were measured using 1D pulse-acquire experiments with samples in the paramagnetic and diamagnetic state, the latter by addition of 2.5 mM (RNC) or 100 x molar excess (isolated protein) sodium ascorbate. For PRE measurements of RNCs, spin-labelled samples were prepared and split into two halves; one half was immediately recorded by NMR at 298 K to produce the paramagnetic spectrum, while the other half was incubated at 277 K. Once the paramagnetic spectrum was acquired (typically 16-24 h), the latter half sample was reduced in sodium ascorbate and the diamagnetic spectrum acquired at 298 K (typically also 16-24 h). To improve signal-to-noise, 5-6 separate RNC samples were prepared and the datasets summed together (using only data corresponding to intact samples) to produce the final spectra. The spectra were fitted to Lorentzian lineshapes, as previously described^2^, which provide accurate measurements of transverse relaxation rates (*R*_2_) and identical to those obtained from direct ^19^F *R*_2_ measurements^3^. Line broadening resulting from the PRE effect resulted in highly overlapped I2 and native state resonances, which were deconvolved by restraining the lineshape fit of the native state using the expected PRE rate bounds (**Supplementary Note 4**).

### AlphaFold structure predictions

AlphaFold3 (ref.^41^, https://alphafoldserver.com) was used to predict structures of FLN5 bihistidine mutants in complex with a metal ion, using Co(II) to replace Ni(II) that is otherwise unavailable. AlphaFold3 was also used to predict the structure of flexible regions of FLN4 that are absent in its crystal structure (PDB: 1WLH). Default settings were applied.

### Cross-linking experiments

Intermolecular cross-links were introduced into FLN5+47 RNC using a similar approach as described for PRE spin-labelling^3^. Samples were reduced overnight at 277 K in Tico supplemented with 2 mM TCEP. TCEP was then removed by buffer exchange into labelling buffer (50 mM HEPES, 12 mM MgCl_2_, 20 mM NH_4_Cl, 1 mM EDTA, pH 8.0). The RNC sample was then incubated with variable concentrations of the cross-linker 1,8-bismaleimido-diethyleneglycol (BM(PEG)_2_) at 298 K, and the reaction quenched with DTT. Cross-linking was assessed by a mass increase of ∼16 kDa (i.e. the mass of uL24) by western blot.

### Proline free energy calculations

We used well-tempered metadynamics^62^ (WT-METAD) simulations to calculate the free energy landscapes capturing the isomerisation state of FLN5 proline P742. All-atom simulations were run using the DES-Amber force field and TIP4P-D water model^63,64^. The calculations were prepared and performed with GROMACS (version 2020)^65^ and PLUMED (version 2.6)^66,67^. Three different models were used to investigate the effect of local structure on P742 isomerisation energetics: an unfolded peptide with <u>P742</u> (underlined) in the middle of the sequence corresponding to FLN5 residues N736-I748 (Ac-NPIKNM<u>P</u>IDVKCI-NMe), a C-terminally truncated FLN5 construct, FLN5 Δ6, populating a folded and disordered C-terminus at equilibrium^11^ and full-length, folded FLN5 (PDB 1QFH, residues K646-G750)^68^. Simulations of the unfolded peptide were initiated from a linear chain built in PyMOL 2.3 (Schrödinger, LLC). All systems were initially centred in a dodecahedral simulation box at least 1.2 nm away from the box edge, solvated and neutralised with MgCl_2_. Energy minimisation of the solvated systems was performed using the steepest descent algorithm using a tolerance of < 1000 kJ mol^-1^ nm^-1^. Equilibration was then performed for 1 ns in the NVT and 1 ns in the NPT ensemble, where the temperature (298 K) was maintained using velocity rescaling^69^ with a coupling constant of 0.1 ps and the pressure (1 bar) was regulated with the Berendsen barostat^70^ a compressibility of 4.5×10^-5^ bar^-1^ and coupling constant of 2 ps. During equilibration all protein heavy atoms were position restrained using a force constant of 1000 kJ mol^-1^ nm^-2^. Position restraints were then removed and pressure coupling switched to the Parrinello-Rahman method^71^ with the same compressibility and coupling constant. A time step of 2 fs was used together with the leap-frog integrator and the LINCS algorithm^72^ to constrain all bonds involving hydrogen atoms. Production simulations with WT-METAD were run in triplicates from different initial velocities for 1 μs each. Van der Waals interactions were computed up to a cut-off distance of 0.9 nm, and short-range electrostatics up to 1 nm. Long-range electrostatics were calculated using the Particle Mesh Ewald (PME) method^73^ with cubic interpolation and a grid spacing of 0.125 nm.

As collective variables for WT-METAD we used two dihedral angles, ζ (Cα_i-1_ – O_i-1_ – C8_I_ – Cα_i_) and Ψ (N_i_ – Cα_i_ – C_i_ – N_i+1_), where *i* corresponds to P742. ζ describes the isomerisation state (∼ 0 for cis and ∼± π for trans in radians) and Ψ describes the the C-terminal amide orientation. Previous work has shown this combination to be efficient for simulating proline isomerisation with WT-METAD^74^. Gaussians were deposited every 1 ps with a width of 0.07 rad for each dihedral angle. The Gaussian height and bias factor were set to 0.8 kJ mol^-1^ and 10, respectively. We used the method described by Tiwary *et al.* to reweight all snapshots to correct for the WT-METAD bias potential (saved every 5 ps)^75^, discarding the first 200 ns (20%) of each trajectory. All ensemble properties including cis/trans populations were then calculated using these weights, with the mean and standard error of the mean calculated from the three independent WT-METAD simulations initiated with different starting velocities. For full-length FLN5 and FLN5 Δ6, we used a collective variable to quantify the fraction of native contacts (Q) formed by the two downstream residues I743 and D744 to correlate C-terminal structure with P742 isomerisation. SMOG2^76^ was used to obtain a list of native heavy atom contacts and Q was calculated using the following switching function^77^:

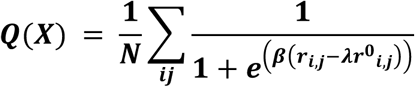

The distance between two heavy atoms is defined as *r_i,j_*, whereas *r*^0^ corresponds to the reference distance. *Δ* and 11 were set to 50 nm^-1^ and 1.8, respectively. WT-METAD simulations of full-length FLN5 were restrained to have a C-terminal Q (CTERM) of > 0.8 using a lower wall restraint in PLUMED (force constant = 10,000 kJ mol^-1^) to maintain the native structure of FLN5 during the simulations.

### All-atom simulations of protein folding

We used ratchet-and-pawl MD (rMD) simulations^78–80^ to sample folding transition paths starting from unfolded conformations. For isolated FLN5, we used SMOG2^76^ to generate a simple structure-based model without any contacts (i.e., no nonbonded attractive potentials) and unfold FLN5. 50 random unfolded conformations were then picked for rMD simulations. Using the DES-Amber force field and TIP4P-D water model^63,64,81^, these structures were then energy minimised and equilibrated using an equivalent protocol as detailed above at 350 K. For the FLN5+47 RNC, we performed thermal unfolding simulations starting from 50 different conformations of folded FLN5+47 on the ribosome obtained in previous work^23^. The RNC was simulated with using the DES-Amber force field and TIP4P-D water model^63,64,81^, because we have shown that this force field results in good agreement of folded FLN5 simulations with NMR data^23^. For thermal unfolding and subsequent relaxation runs, we used hydrogen mass repartitioning^82^ and a 4 fs timestep with LINCS. GROMACS 2020 and PLUMED 2.6 were used for these simulations. Temperature and pressure coupling settings were identical to the production runs used for the proline free energy calculations described above. All ribosome heavy atoms (and C-terminal nascent chain residue) were position restrained at all times using a force constant of 1000 kJ mol^-1^ nm^-1^. Unfolding was performed at 400 K and 1 bar for 25 ns in the presence of lower wall restraints on the radius gyration calculated from residues 646-750 Cα atoms at 3 nm with a force constant of 10,000 kJ mol^-1^. The total fraction of native contacts, as defined in the previous section, was restrained to be below 0.02 using a force constant of 10,000 kJ mol^-1^. To ensure all starting structures had a trans peptide bond for P742 (P742 is in the cis conformation in the native structure), we additionally restrained the absolute value of the ζ dihedral angle to be above 2.62 rad (∼150°). These unfolded structures were then relaxed at 350 K and 1 bar for 1 ns in the absence of wall restraints (but with ribosome position restraints).

Enhanced sampling is achieved with the rMD method (adiabatic-bias MD, ABMD, in PLUMED). We used GROMACS 2020 and PLUMED 2.6 for rMD simulations. In this approach, simulations proceed in the absence of a biasing potential when spontaneous progression towards a pre-defined target state (i.e., the folded state of FLN5) occurs. However, a soft biasing potential is applied when the system moves in the opposite direction^78,79^:

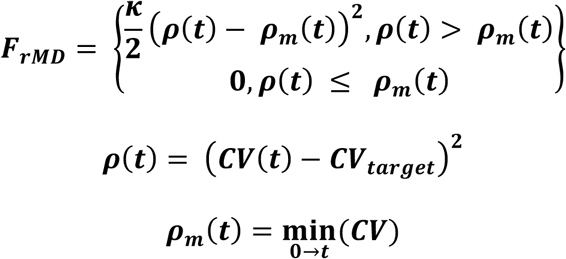

The force constant, *K*, controls the magnitude of the bias potential, *CV(t)* is the instantaneous value of a defined collective variable at time, *t*, and *CV_target_* is its value in the target (folded) state. rMD therefore effectively promotes transitions from an initial (unfolded) state to the final (native state) using a reaction coordinate. In our rMD simulations, we set *K* and *CV_target_* to 100 kJ mol^-1^ and 0, respectively. As a collective variable, we used the difference contact map between a given conformation and the native state^78,79^, defined by heavy atom native contacts, *j,* within a 0.6 nm distance cut-off obtained the shadow algorithm implemented in SMOG2^76^:

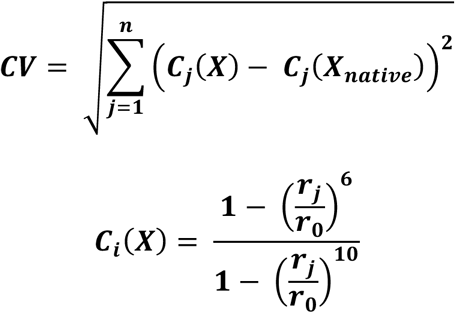

Here, *r_j_* is the native contact distance for atom pair *j* and *r_0_* was set to 0.5 nm. For FLN5, this approach includes 1,009 heavy atom contacts calculated using the energy minimised crystal structure of FLN5. For all 50 starting structures, we ran 36 independent rMD simulations starting from randomly assigned initial velocities for a time of 5 ns, totalling 1800 trajectories and 9 µs of biased sampling. Simulations were run in the NPT ensemble at 350 K, using the same temperature and pressure coupling parameters as described above. The integration time step was 2 fs. For the RNC, position restraints on the heavy atoms of the ribosome and C-terminal NC residue at the peptidyl transferase centre were kept at 1000 kJ mol^-1^ nm^-1^. We then calculated the RMSD over Cα atoms of residues 646-750 for all trajectories with respect to the energy minimised crystal structure of FLN5. Trajectories reaching an RMSD of less than 0.2 nm were defined as successful folding transition paths, resulting in 321 and 226 successful folding transitions for isolated FLN5 and FLN5+47 RNC, respectively. These trajectories were analysed further and we calculated the total fraction of native contacts and native contacts between specific structural elements of the protein to project the transition path landscape onto these variables to investigate the folding pathway and intermediate states.

### Unbiased all-atom simulations of coTF intermediates

To select starting structures of the I1 coTF intermediate, we considered all potential structures belonging to the local free energy minimum observed in rMD simulations between a fraction of a global Q of 0.55 < Q < 0.68 (**Figure S5b-f**). Given this intermediate free energy minimum corresponds to structures with a folded G-strand (**Figure S5f**) and our proline free energy calculations show that P742 is in the cis configuration in this case (**Figure S4**), we only selected structures with P742 in cis. To ensure the selection of physically reasonable structures, we sorted the structures by ascending distance RMSD (dRMSD, relative to the energy-minimised native state crystal structure) of all heavy atoms belonging to residues F691-G750. These residues correspond to the natively folded region of this intermediate state predicted by rMD (**Figure S5c**). The top 100 structures were then clustered using the GROMOS algorithm^83^ implemented in gmx cluster, using all-atom RMSD of the entire nascent chain without alignment and a cut-off of 0.9 nm to select structures with different orientations on the ribosome. The top 5 cluster centroids (**Figure S5g**) were then used for unbiased MD simulations.

Since rMD simulations only sampled one intermediate (I1, **Figure S5**) and due to the structural similarity between the I2 coTF intermediate and an intermediate previously characterised for isolated FLN5 Δ6^3,11,38^, we randomly selected five structures from the lowest free energy cluster of the NMR-derived structural ensemble of isolated FLN5 Δ6^11^. We then built the C-terminal residues of FLN5 in an extended conformation and joined these models with the NC linker for FLN5+47 used in previous work^23^. Unbiased MD simulations were then initiated from these structural models.

For all input structures of I1 and I2, we placed these complexes in the centre of a dodecahedron box and added solvent atoms and MgCl_2_ to neutralise the charge of the system. The box size was determined by gmx editconf using a minimum distance of 3 and 2 nm from the box edge for I1 and I2, respectively. This resulted in system sizes of 1.2 – 1.7 million atoms. After energy minimisation, we first gently heated the simulation boxes from 0 to 298 K over a 500 ps using a time step of 0.5 fs and LINCS constraints for all bonds involving hydrogen atoms. The systems were then equilibrated at 298 K in the NVT ensemble (velocity rescaling thermostat with a coupling constant of 0.1 ps) for 500 ps using a 2 fs time step, followed by a 1 ns equilibration step in the NPT ensemble at 1 bar (Berendsen barostat with a time constant of 2 ps). During the equilibration protocol, all RNA and protein heavy atoms were position restrained (1000 kJ mol^-1^ nm^-2^). These restraints were then removed for all NC atoms (except the C-terminal residue at the peptidyl transferase centre) for a further 1 ns equilibration simulation (NPT, Parrinello-Rahman barostat, 2 ps coupling constant) and production simulations. We performed three unbiased simulations starting from different initial velocities for each starting structure lasting 1 μs, resulting in a total of 15 μs for each coTF intermediate state.

RMSF and SASA analyses were performed with gmx rmsf and sasa^65^, respectively. Alignment prior to RMSF calculation was done using Cα atoms of residues in the folded region of the intermediate state, F691-G750 (C-strand to C-terminus) and K646-L733 (N-terminus to F-strand) for I1 and I2, respectively. The fraction of native contacts (Q) was calculated using PLUMED as described above. The radius of gyration was computed using all Cα atoms of the FLN5 domain (residues K646-G750) with PLUMED to compare the compactness of the coTF intermediates with the native state. Contact maps, ribosome interactions, distances, and secondary structure content were calculated using the MDTraj^84^ and MDAnalysis^85^ Python libraries. The rotational correlation times of the coTF intermediates were calculated as previously described^23^. The rotation of the domain was defined using backbone heavy atoms belonging to the same regions as defined for the RMSF analysis of I1 and I2. Structural analyses were compared with the native state ensemble of FLN5 previously obtained^23^ with the DES-Amber force field. All ensemble properties were averaged over the entire 15 μs ensemble and errors are reported as the standard error of the mean obtained from the 15 independent 1 μs trajectories.

The structural ensembles were clustered using for the purpose of structural visualisation using gmx cluster^65^ and the GROMOS algorithm^83^. Alignment and RMSD calculations for clustering were performed using the same residue ranges as for the RMSF analysis and all backbone heavy atoms. A similarity cutoff of 0.2 and 0.1 nm were chosen for I1 and I2, respectively, due to these values resulting in similar cluster populations (**Table S4**). Structures from the most populated cluster were then used for visualisation in the main text figure (**Figure 3a-d**).

### Simulations of fluorinated protein variants in isolation

We used the CHARMM36m^86^ and ff15ipq^87,88^ force fields to simulate the following isolated FLN5 variants labelled with 4-trifluoromethyl-L-phenylalanine (tfmF): 655tfmF, 691tfmF, 705tfmF, 715tfmF, 719tfmF, 727tfmF, 732tfmF, and 740tfmF. We used an identical simulation protocol as described previously^39^. Production simulations were run for 1 μs, saving coordinates for analysis every 0.1 ns. Titin I27 variants lacking the A-strand (residues 1-7 deleted) were simulated with the ff15ipq^87^ force field for 10 μs in triplicates to sufficiently sample the folding intermediate. GROMACS tools gmx sasa, rmsd, and rmsf were used for general analyses of the CF_3_ group (SASA) and protein structure/dynamics (rmsd and rmsf). Hydrogen bonds between fluorine (acceptor) and donor atoms were analysed with gmx hbond. To estimate hydrogen bond lifetimes, we calculated the area under the averaged autocorrelation function from all hydrogen bond existence functions^89,90^. Distances were calculated with in-house Python scripts using MDAnalysis^85^, and CF_3_-aromatic interactions were analysed with an open-source Python script (https://github.com/julian-streit/RingCurrents19F).

Simulations isolated FLN5 fragments mimicking coTF intermediates I1 and I2 were performed using the ff15ipq force field^87^ and minimal constructs comprising the folded core of each intermediate. We used residues Ac-A683-G750-NMe and S645-D744 for I1 and I2, respectively. Three different starting structures were taken from the same rMD-generated models as used for the unbiased RNC simulations (I1) and FLN5 Δ6 structural ensemble^11^ (I2). Independent simulations were performed using the same protocol as for full-length fluorinated proteins from these starting structures with random initial velocities to obtain three replicates where the I1/I2 topology was maintained during the 1 μs simulations. This was assessed by having at least one ^19^F NMR residue pair with a native-like distance per Δ-strand pair (within 0.1 nm of the average distance calculated for the FLN5+47 native state^23^. The native β-strand pairs measured by ^19^F NMR for I1 were CF/FG, and AB/BE/DE/CF for I2.

### 19F PRE distance calculations

^19^F NMR PRE rates *ρ_2_* were converted to distances, *r*, using the Solomon-Bloembergen equation^91^, where *ω_F_* and *τ_C_* are the Larmor frequency of fluorine-19 and rotational correlation time, respectively:

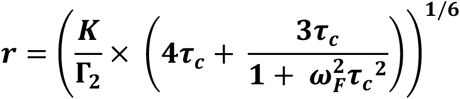

*K* (1.092 x 10^-44^ m^6^s^-2^) is defined as a product of physical constants including the vacuum permeability, *μ_0_*, gyromagnetic ratio of fluorine-19, *ψ_F_*, electron g-factor, *g*, Bohr magneton, *μ_B_*, and the spin number of fluorine-19, S.

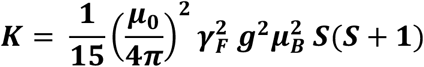

Values of*τ_C_* have been determined in previous studies^37^, and for RNCs were determined from the linewidths using an empirical relationship recently determined^3^. To extrapolate the expected PRE of native FLN5 on the ribosome, we therefore used the measured PRE distance in isolation and RNC-specific *τ_C_*:

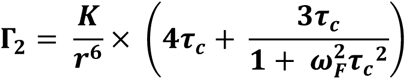

We fitted the ribosome model used in the MD simulations to each cryo-EM map representing the four FLN5+47 dataset classes (AP1, AP2, AP5 and P5) (Mitropoulou, et al, in preparation). The obtained rotation-translation matrix was applied in Python using the MDAnalysis library to fit the NC structure from each frame of the MD trajectory (native, I2 and I1 states) into the NC density. Next, we used scripts from cryoENsemble^43^ to generate synthetic cryo-EM maps for each MD frame at 8 Å resolution. The correlation coefficient between each synthetic map and the corresponding experimental cryo-EM map was then calculated following the definition of map correlation by ChimeraX^92^.

As an alternative approach, we also used the cryo-EM maps to reweight the combined trajectories corresponding to native, I2 and I1 states using the cryoENsemble iterative strategy^43^. Prior to reweighting we clustered each trajectory using gromos algorithm from the gmx cluster, using a 1-nm cut-off. The obtained clustered trajectory, comprising 2,294 conformations (1,091 from I1, 706 from I2 and 497 from native state), was used in four cryoENsemble reweighting runs in each of the cryo-EM maps of FLN5+47, following the previously described protocol^43^.

## Data availability

The structural ensembles of FLN5+47 I1 and I2, and NMR data will be deposited on Zenodo upon publication. This study made use of the following public datasets deposited in the PDB (https://wwww.rcsb.org): 1QFH, 1TIT, 1WLH, and 2W0P.

## Code availability

Pulse sequences used for the acquisition of NMR data, and Matlab and Julia codes used to fit the ^19^F NMR data are available on Github (https://github.com/shschan/). Python scripts used to predict aromatic ring interactions for fluorinated protein variants are available on Github (https://github.com/julian-streit/RingCurrents19F).

## Acknowledgements

This work was supported by a Wellcome Trust Investigator Award (to J.C., 206409/Z/17/Z).

J.O.S. was supported by a BBSRC London Interdisciplinary Biosciences Doctoral Programme studentship. We acknowledge use of the UCL Biomolecular NMR Centre. This project made use of time on HPC resources on Archer2 (ARCHER2 UK National Supercomputing service, https://www.archer2.ac.uk) granted via the UK High-End Computing Consortium for Biomolecular Simulation, HECBioSim (https://www.hecbiosim.ac.uk), supported by EPSRC (grant no. EP/R029407/1 and EP/X035603/1). We also acknowledge the EuroHPC Joint Undertaking for awarding this project access to the EuroHPC supercomputer LUMI, hosted by CSC (Finland) and the LUMI consortium through a EuroHPC Regular Access call and the Baskerville Tier 2 HPC service (https://www.baskerville.ac.uk/). Baskerville was funded by the EPSRC and UKRI through the World Class Labs scheme (EP/T022221/1) and the Digital Research Infrastructure programme (EP/W032244/1) and is operated by Advanced Research Computing at the University of Birmingham. We additionally acknowledge the use of the UCL Myriad, Kathleen, and Young High Performance Computing Facility, and associated support services, in the completion of this work.

## Author contributions

S.H.S.C., J.O.S., and J.C. designed and conceptualised the project. S.H.S.C. and J.O.S. produced the samples and performed the NMR experiments. S.H.S.C. analysed the NMR experiments. J.O.S. performed and analysed the MD simulations. T.W. and A.M. performed the cryo-EM analyses. A.M.E.C. and I.V.B. contributed to the initial C740 PRE experiments. S.H.S.C., J.O.S., L.D.C., and J.C. sourced the computational resources and acquired the funding. S.H.S.C. and J.C. supervised the project. S.H.S.C., J.O.S., and J.C. administered the project. S.H.S.C. wrote the original draft, S.H.S.C., J.O.S., and J.C. edited the manuscript, and all authors reviewed the manuscript.

