## Supplementary Information for "Structures of protein folding intermediates on the ribosome"

### Supplementary Note 1: $^{19}$F NMR of intermediate states on the ribosome

The presence and number of broad $^{19}$F NMR resonances attributable to intermediate states was confirmed through a series of analyses and additional experiments. In addition to quantitative lineshape analyses in the frequency domain (**Figure 1h**, **S1a**), the number of peaks fitted in each spectrum was confirmed by statistical tests of fits performed on data analysed in the time domain[2] (**Methods, Table S3**). The RNC spectra were additionally compared against those measured for RNCs of a fully unfolded, folding-incompetent mutant (FLN5+47 Tyr719Glu) labelled at the same site (**Figure S1a**), which showed a narrower linewidth for the unfolded state (compared to the putative intermediate state resonances, **Table S1**). We also incubated FLN5+47 tfmF718 671Phe RNCs in mild denaturant (2.5 M urea, **Figure S1a**). Under these conditions, the folding equilibrium is expected to be shifted, and indeed, four NMR peaks were observed with the additional resonance attributable to the unfolded state, having a narrow linewidth (~11 Hz) and a random coil chemical shift (**Figure S1a**). The remaining resonances showed reduced integrals (i.e. populations) and line broadening as expected (**Tables S1-2**), the latter resulting from reduced interactions between the nascent protein and ribosome surface in urea, and which improved resolution between the overlapped resonances. Finally, the two broad resonances were also found at a shorter RNC length, with broader linewidths resulting from increased ribosome surface interactions closer to the ribosome as expected[5] (**Figure S1a**). Similar analyses and experiments were performed for the remaining $^{19}$F-label sites (**Figure S1a, Tables S1-3**). Collectively, the data and analyses confirm the presence of broad resonances attributable to two distinct intermediate states across the $^{19}$F NMR spectra of RNCs.

### Supplementary Note 2: Ratchet-and-pawl simulations and starting structures of unbiased MD simulations

We used all-atom MD simulations as an orthogonal means to assess the coTF intermediate structures and to obtain starting structures for subsequent unbiased MD simulations. We initially performed ratchet-and-pawl (rMD) simulations[78,79] using the DES-Amber force-field in explicit solvent[63,64,81] (**Methods**) to model the sequence of folding events from an unfolded FLN5+47 RNC to a native-like conformation (**Supplementary Videos 1-2**, **Figure S5**).

Analysis of the folding trajectories showed a largely hierarchical folding pathway, where β-strand pairs are progressively formed (**Figure S5b-d**). Crucially, we identified a local minimum within the kinetic energy landscape, corresponding to an intermediate state. The structural ensemble possessed a native-like core comprising the C- to G-strands, and was found on-pathway and prior to the folding of the remaining N-terminal A-B strands to form the fully native state (**Figure S5b-d**). Additional calculations show that native G-strand folding is thermodynamically dependent on the isomerisation state of the conserved natively cis-Pro742 (**Figure S4**); within this intermediate, Pro742 is therefore expected to adopt a cis-conformation. These structural features appeared to align most closely with NMR data of the I1 state (**Figure 2a-b**, and ref.[2]), and we therefore assessed this intermediate state further by sampling the structures and dynamics from the folding trajectories in additional MD simulations (**Figure 3, S6**).

In contrast, we could not identify a local energy minimum corresponding to a potential I2 state (**Figure S5b-f**), likely due to methodological limitations of the rMD approach. However, given its structural similarity to a folding intermediate populated by an isolated, truncated variant of FLN5[11] (**Figure 2c-d**), as also found from previous coarse-grained structure-based models[2], we instead used previously determined NMR structures of the isolated intermediate[11], having a disordered G-strand and Pro742 in the trans conformation[11] (**Figure S4**) as starting models of MD simulations. We tethered the isolated intermediate to the ribosome ensuring the same sequence as FLN5+47 (**Methods**). Extensive sampling of the structure and dynamics of the I1 and I2 models were subsequently performed using unbiased all-atom MD simulations (**Figure 3, S6, Supplementary Videos 3-4**)

**Supplementary Note 3: Assessing intermediate structure distance distributions against chemical shift data**

To assess our structural ensembles, we calculated distance distributions between pairs of residues that were used for $^{19}$F-labelling (typically $C_\beta$-$C_\beta$ atomic distances), and compared these analyses to both those of the native structure[23] and also the experimental chemical shifts, as the latter are engineered to probe the inter-residue contact/distance of the label pair (**Figures 1f-h**, **S1a**). This simple analysis rationalised almost all of the observed chemical shifts.

More accurate comparisons can be made by explicitly modelling the $^{19}$F-label pair (tfmF and the accompanying aromatic residue) and calculating the predicted geometric factor $(1-3\cos^2\theta)/r^3$, where $r$ and $\theta$ are the distance and angle of the $CF_3$ group in tfmF relative to the aromatic ring. We first modelled the native and I1 intermediate state with $^{19}$F-label pairs across the F-G strands. Both label pairs (726tfmF 746Phe and 728tfmF 744His) yielded similar geometric factors for the I1 and native states (**Figure S6k**), as also reflected by their inter-residue $C_\beta$-$C_\beta$ distances measured without explicit $^{19}$F-label pairs (**Figure 3h**) and by the experimental data (chemical shifts and bihistidine metal binding, **Figures 1-2**). In contrast, assessing the D-E strand contacts (710tfmF 716His and 718tfmF 706His), we found substantially reduced (>50%) geometric factors for I1 relative to the native state (**Figure S6k**) for both labelling sites, but native-like $C_\beta$-$C_\beta$ ($C_\alpha$ for Gly716) distances calculated for their structures without explicit $^{19}$F-label pairs (**Figure 3h**). These calculations indicate that the D-E strands form native-like secondary structures in I1, with more subtle sidechain conformational changes that result in reduced ring current interactions (relative to native state), and accounting for their random coil chemical shifts.

For natively buried probes (727tfmF, 715tfmF) whose chemical shifts report on specific side chain-amide backbone contacts (**Figure S2**), we calculated hydrogen bond propensities (**Figure S6l-m**) and found that native interactions remained in I2 structures, but not in I1, reflective of their extent of structure formation and indeed their chemical shifts.

**Supplementary Note 4: $^{19}$F PRE measurements as validation of structural models**

The paramagnetic effect results in substantial line broadening in the $^{19}$F NMR spectrum of the RNC (**Figure 4c, S7b**), as expected, and indicative of Ni(II) binding to the nascent protein as designed. While it is evident from the raw spectra that there is only a minimal PRE effect on the I1 resonance, with a significantly greater effect on the I2 state, the increased line broadening results in severe overlap of resonances, complicating quantitative lineshape analyses. We note that this could not be alleviated by improving the signal-to-noise; indeed, the paramagnetic spectra of the RNCs are each summed from 5-6 separate samples (~100 h total acquisition time per PRE-label). Restraints were therefore required to fit lineshapes to the paramagnetic spectrum. We calculated the expected PRE rate of the native state in

FLN5+47, using the experimental PRE rate measured for the isolated, natively folded protein (**Figure S7a**) and scaled this with the rotational correlation time on the RNC, since the native structure is the same on/off the ribosome[23,37]. The rotational correlation time was determined using linewidths measured from the diamagnetic spectra (**Figure S7c**) and a previously empirically determined correlation between these two parameters[3,37]. The expected PRE rate was then converted to a linewidth (with propagated errors) of the native state resonance in the paramagnetic spectrum, permitting free fits of the I1 and I2 lineshapes (**Figures 4c, S7b**). We then converted the measured linewidths of I1 and I2 to a PRE rate (**Figure S8d**) and distance (**Figure 4d**). Distances were calculated using the rotational correlation time (measured in the same manner as for the native state), and assuming that the interaction vector in the molecular frame is the same across I1, I2, and the native state (**Methods**). From these calculations, we find the distance between 655tfmF and 673Cys-MTSL in I2 to be native-like, as expected from the 655tfmF chemical shift (probing Phe675, **Figure 1f**), validating our approach.

We compared the experimental distances to those calculated from the determined structural ensembles. As the rotamer distribution of the flexible MTSL spin label cannot be accurately determined (even when explicitly modelled), we instead calculated the distance between OH of 655tfmF and the $C_\beta$ of the residue bearing the spin label. Comparison between the experimental and calculated distances for the native state (**Figure 4d**) indicate a small systematic (scaled) discrepancy, with the calculated distance being slightly underestimated for 675Cys-MTSL label, but overall good agreement, showing greater distances for I1 (relative to native) but native-like distances for I2.

**Supplementary Note 5: Cross-linking experiments and cryo-EM fits**

We performed cross-linking experiments to examine specific ribosome interactions. Our structures showed that all states (I1, I2, N) do not substantially contact uL24 via the N-terminal residue Ala648 (<5% within 2 nm**, Figure S8d**); accordingly, we did not detect cross-links between FLN5+47 RNC Ala648Cys and uL24 (Gly90Cys) on CRISPR-modified ribosomes (**Figure S8e**). In contrast, cross-links between uL24 (Gly90Cys) and FLN5+47 RNC labelled at Asp699Cys were found, and could be rationalised by the increased contact (~8% within 2 nm) as observed in the I1 structures (**Figure S8d-e**).

We have previously obtained cryo-EM maps of FLN5+47 and fitted natively folded FLN5 conformations to the densities (Mitropoulou *et al.*, in preparation). However, slow chemical exchange of the native state with both I1 and I2 (ref.[2]) indicates that each conformation likely contributes to the observed cryo-EM densities. Therefore, we firstly compared cross-correlations between single structures within our three structural ensembles (native[23], and I1 and I2 states) with the maps; high cross-correlations were obtained when fitting the intermediate state structures, which were similar or better to those obtained using native state conformations (**Figure S8a**), supporting the validity of our models. In an orthogonal approach, we used the cryo-EM maps to reweight the structural ensembles of I1, I2, and N using cryoENsemble[43]. Cross-correlations improved when combining all three ensembles together for reweighting (**Figure S8b-c**), compared to individual ensembles (**Figure S8c**), indicating that the densities most likely derive from heterogenous populations of RNCs of different conformational states, therefore crucially showing that addition of our I1 and I2 models improved the overall fit.

Altogether, the biochemical data and cryo-EM densities support the determined ensembles of intermediate structures, their ribosome interaction sites, and orientations.

**Supplementary Note 6: Intermediate structures of I27**

We performed all-atom MD simulations to model structures of an isolated I27 intermediate analogue[44]. The intermediate structure comprises a well-folded A'-G core with a detached A-strand, as expected[44,45] (**Figure S9d**), permitting closer interactions between A'-G strands; this is reflected by quantitatively stronger ring current interactions between 14tfmF and 87His (relative to native state, **Figure S9e**), and which therefore rationalises the observed highly shielded $^{19}$F chemical shift of the isolated and ribosome-bound intermediate (**Figures 5d, S9c**). Indeed, agreement between another $^{19}$F chemical shift of I27 RNCs with calculated ring current contacts across the isolated intermediate models (59tfmF, **Figure S9**) suggest that the same intermediate structure is populated on the ribosome. Meanwhile, the chemical shifts of the remaining intermediate (I1) state resonances (**Figure S9c**) indicate a less ordered structure, although overall maintaining a native-like hydrophobic core[3] (**Figure S9c**), with inter-strand contacts identical to those observed in FLN5 I1 and thus suggestive of a similar structure.

**Supplementary Video 1**

Exemplar rMD trajectory of FLN5+47 RNC showing folding sequence via I1 conformation with disordered A-B strands.

**Supplementary Video 2**

Exemplar rMD trajectory of FLN5+47 RNC showing folding sequence via I1 conformation with a detached A-B β-hairpin.

**Supplementary Video 3**

Exemplar unbiased MD trajectory (1 μs) of the FLN5+47 RNC I1 state with a detached A-B β-hairpin.

**Supplementary Video 4**

Exemplar unbiased MD trajectory (1 μs) of the FLN5+47 RNC I2 state with a detached G-strand.

**Supplementary Figures**

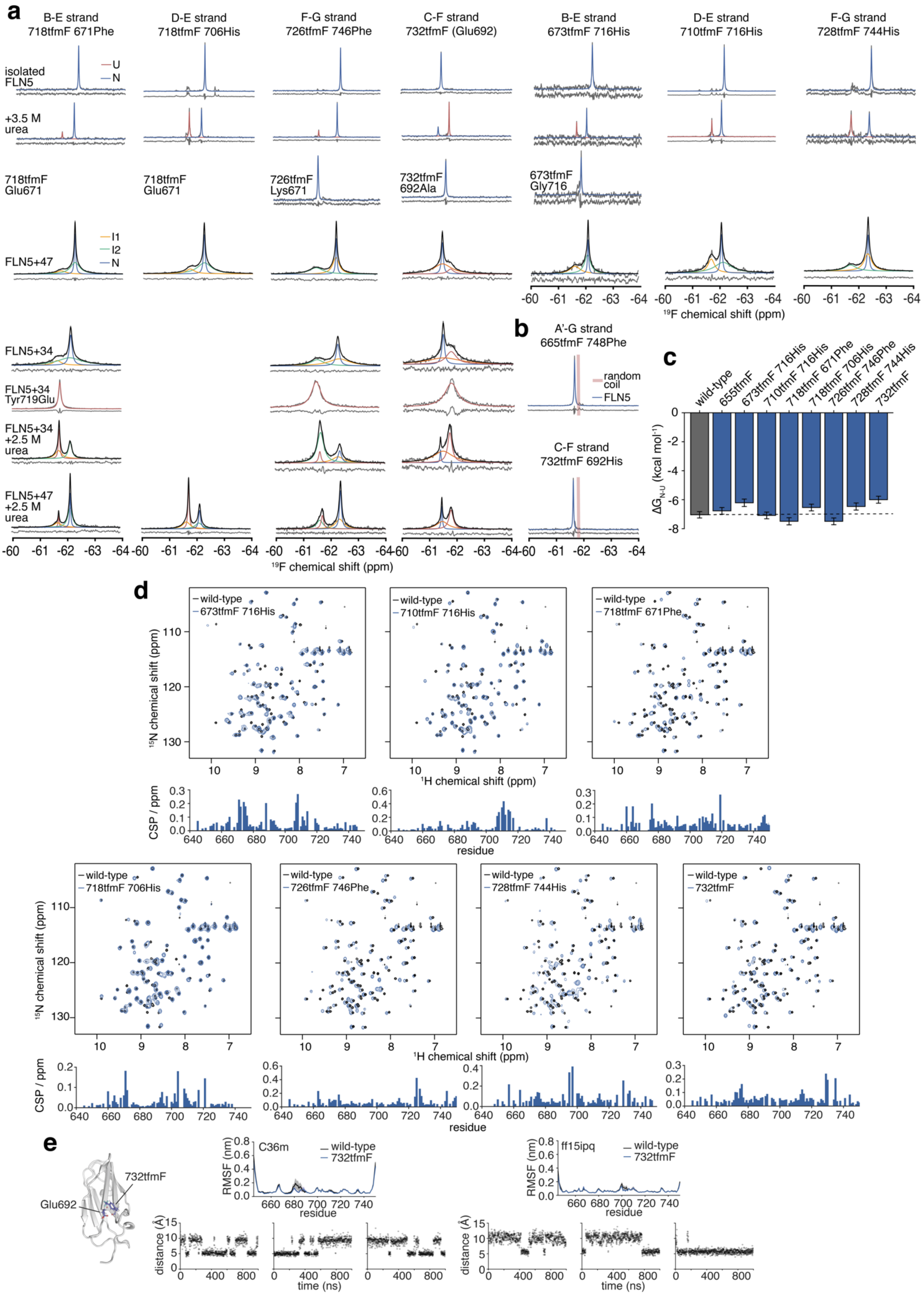
a
B-E strand 718tfmF 671Phe
D-E strand 718tfmF 706His
F-G strand 726tfmF 746Phe
C-F strand 732tfmF (Glu692)
B-E strand 673tfmF 716His
D-E strand 710tfmF 716His
F-G strand 728tfmF 744His
isolated FLN5
U
N
+3.5 M urea
718tfmF Glu671
718tfmF Glu671
726tfmF Lys671
732tfmF 692Ala
673tfmF Gly716
FLN5+47
I1
I2
N
FLN5+34
FLN5+34 Tyr719Glu
FLN5+34 +2.5 M urea
FLN5+47 +2.5 M urea
19F chemical shift (ppm)
b
A'-G strand 665tfmF 748Phe
random coil
FLN5
C-F strand 732tfmF 692His
c
wild-type
655tfmF
673tfmF 716His
710tfmF 716His
718tfmF 671Phe
718tfmF 706His
726tfmF 746Phe
728tfmF 744His
732tfmF
ΔGN-U (kcal mol-1)
d
wild-type
673tfmF 716His
710tfmF 716His
718tfmF 671Phe
718tfmF 706His
726tfmF 746Phe
728tfmF 744His
732tfmF
15N chemical shift (ppm)
1H chemical shift (ppm)
CSP / ppm
residue
e
732tfmF
Glu692
C36m
ff15ipq
RMSF (nm)
distance (Å)
time (ns)

**Figure S1 Characterisation of solvent-exposed tfmF-labelled variants of FLN5 on and off the ribosome. a,** $^{19}$F NMR spectra of FLN5 tfmF-labelled at different positions, as isolated protein, with/without urea and with/without the accompanying aromatic label residue, and as RNC. Observed spectra shown in grey were fitted and assigned to their conformational states (coloured). Total fitted spectra are shown in black, and residual spectrum after fitting are shown below. Spectra were recorded at 500 MHz and 298 K. **b,** $^{19}$F NMR spectra of FLN5 tfmF-labelled at different positions. Red vertical line indicates random coil chemical shift range, and therefore the limited chemical shift dispersion of the variants. Observed spectra shown in grey were fitted (blue), with the residual spectrum after fitting shown below. Spectra were recorded at 500 MHz and 298 K. **c**, Gibb's free energies of folding of different tfmF-labelled FLN5, determined by lineshape analyses of spectra shown in a. Errors were determined by bootstrapping of residuals from lineshape fits. Values for wild-type and 655tfmF FLN5 were previously reported[2,11]. **d**, 2D $^{1}$H,$^{15}$N-SOFAST HMQC spectra of tfmF-labelled FLN5 overlaid with that of wild-type FLN5. Chemical shift perturbations (CSPs) are shown below. Spectra were recorded at 500 MHz (wild-type at 800 MHz) and 298 K. **e**, Simulations of FLN5 732tfmF. Left shows a structural model highlighting the 732tfmF and Glu692 sidechains. Top middle/right shows average (± s.e.m.) $C_\alpha$-RMSF of wild-type and 732tfmF FLN5 simulated with C36m and ff15ipq parameters. Bottom middle/right shows distances between the centres of mass of tfmF $CF_3$ and Glu692 carboxylic acid group, for 3 independent simulations using C36m and ff15ipq parameters.

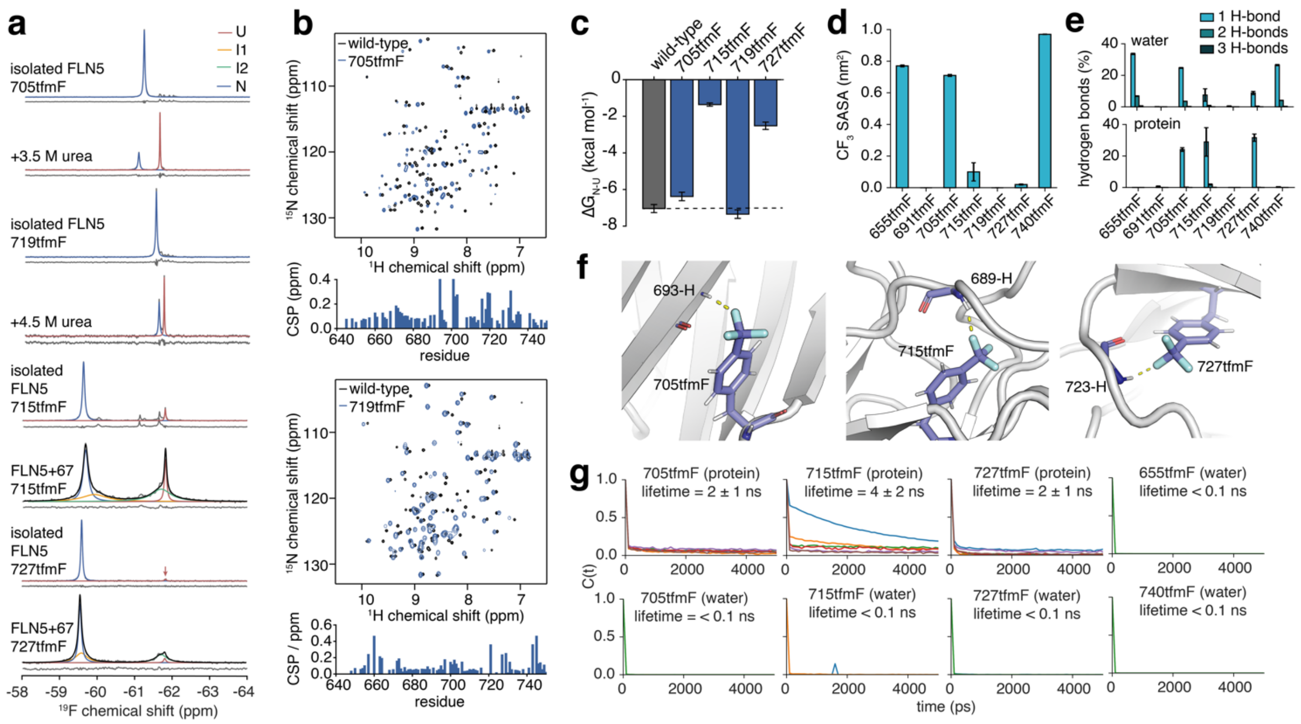


**Figure S2 Characterisation of natively buried tfmF-labelled variants of FLN5 on and off the ribosome. a,** $^{19}$F NMR spectra of FLN5 tfmF-labelled at different positions, as isolated protein, with/without urea and with/without the accompanying aromatic label residue, and as RNC. Observed spectra shown in grey were fitted and assigned to their conformational states (coloured). Total fitted spectra are shown in black, and residual spectrum after fitting are shown below. Spectra were recorded at 500 MHz and 298 K. **b,** 2D $^{1}$H,$^{15}$N-SOFAST HMQC spectra of tfmF-labelled FLN5 overlaid with that of wild-type FLN5. Chemical shift perturbations (CSPs) are shown below. Spectra were recorded at 500 MHz (wild-type at 800 MHz) and 298 K. **c**, Gibb's free energies of folding of different tfmF-labelled FLN5, determined by lineshape analyses of spectra shown in a. Errors were determined by bootstrapping of residuals from lineshape fits. Value for wild-type FLN5 was previously reported[11]. **d-g,** Simulations and interactions of additional FLN5 tfmF sidechains with ff15ipq parameters. Solvent-exposed sites (655tfmF and 740tfmF) were used as controls. For wild-type, 655tfmF and 740tfmF, three independent simulations of 1 μs were performed. For 691tfmF, 705tfmF, 715tfmF, 719tfmF, and 727tfmF six simulations of 1 μs (and in three simulations water coordinates were saved for analysis). **d,** Solvent-accessible surface area of the $CF_3$ group (mean ± s.e.m.) for all variants. **e,** Population of hydrogen bonds (H-bonds) between $CF_3$ and water, and between $CF_3$ and protein atoms, for all variants (mean ± s.e.m.). **f,** Most common hydrogen bond contacts formed by 705tfmF, 715tfmF and 727tfmF with amide groups of residues 693, 689 and 723, respectively. **g,** Averaged hydrogen bond existence autocorrelation functions and estimated lifetimes (mean ± s.e.m. or upper bound, from the area under the curve). Individual lines represent simulations replicates.

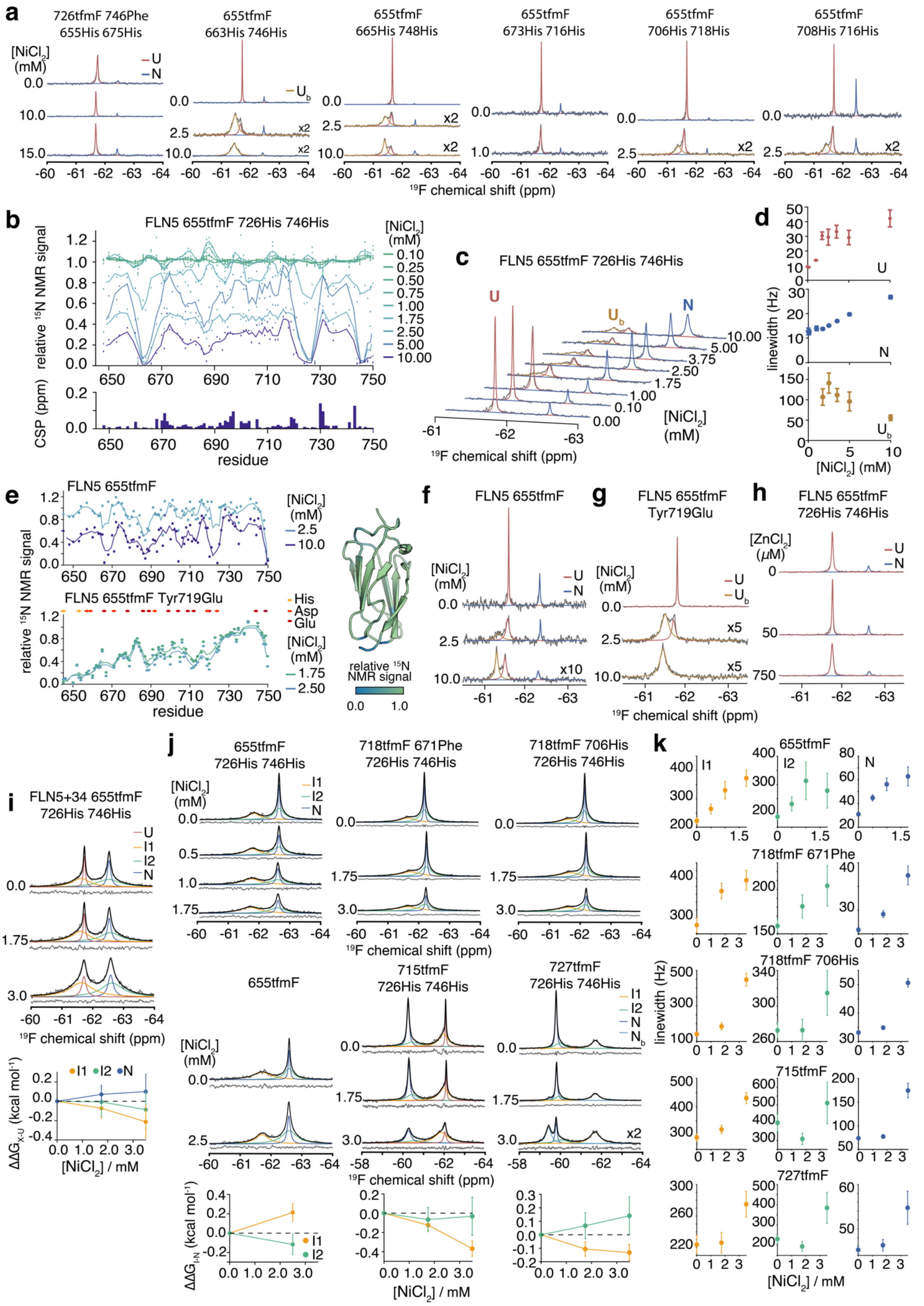
a
726tfmF 746Phe 655His 675His
655tfmF 663His 746His
655tfmF 665His 748His
655tfmF 673His 716His
655tfmF 706His 718His
655tfmF 708His 716His
[NiCl2] (mM)
U
N
Ub
x2
19F chemical shift (ppm)
b
FLN5 655tfmF 726His 746His
relative 15N NMR signal
CSP (ppm)
residue
[NiCl2] (mM)
0.10
0.25
0.50
0.75
1.00
1.75
2.50
5.00
10.00
c
FLN5 655tfmF 726His 746His
19F chemical shift (ppm)
d
linewidth (Hz)
[NiCl2] (mM)
e
FLN5 655tfmF
FLN5 655tfmF Tyr719Glu
His
Asp
Glu
relative 15N NMR signal
residue
relative 15N NMR signal
f
FLN5 655tfmF
g
FLN5 655tfmF Tyr719Glu
h
FLN5 655tfmF 726His 746His
[ZnCl2] (μM)
x5
x10
i
FLN5+34 655tfmF 726His 746His
I1
I2
ΔΔGX-U (kcal mol-1)
[NiCl2] / mM
j
655tfmF 726His 746His
718tfmF 671Phe 726His 746His
718tfmF 706His 726His 746His
655tfmF
715tfmF 726His 746His
727tfmF 726His 746His
Nb
ΔΔGI-N (kcal mol-1)
k
655tfmF
718tfmF 671Phe
718tfmF 706His
715tfmF
727tfmF
linewidth (Hz)
[NiCl2] / mM

**Figure S3 Engineering ligand binding sites to stabilise specific conformational states.** **a,** $^{19}$F NMR spectra of isolated FLN5 with bi-histidine mutations at different positions (across different β-strands) and with varying concentrations of $NiCl_2$. All show no substantial stabilisation of the natively folded state with $NiCl_2$. **b**, Top shows residue-resolved $^{1}$H,$^{15}$N-SOFAST HMQC signal intensities of FLN5 655tfmF 726His 746His in varying $NiCl_2$ concentrations relative to without $NiCl_2$. Bottom shows $^{1}$H,$^{15}$N-chemical shift perturbations of FLN5 655tfmF 726His 746His in 10 relative to 0 mM $NiCl_2$. **c,** $^{19}$F NMR spectra of isolated FLN5 655tfmF 726His 746His with varying concentrations of $NiCl_2$. **d**, Linewidths determined by lineshape analysis of spectra shown in c. Errors were determined by bootstrapping of residuals from lineshape fits. **e,** Residue-resolved $^{1}$H,$^{15}$N-SOFAST HMQC signal intensities of FLN5 655tfmF (without bi-histidine mutations, 2.5 mM dataset mapped onto the FLN5 crystal structure, PDB 1QFH, right) and FLN5 655tfmF Tyr719Glu (the latter unfolded[36]) in varying $NiCl_2$ concentrations relative to without $NiCl_2$. **f,** $^{19}$F NMR spectra of isolated FLN5 (without bi-histidine mutations) in varying $NiCl_2$ concentrations. **g,** $^{19}$F NMR spectra of isolated FLN5 Tyr719Glu (without bi-histidine mutations, and unfolded[36]) in varying $NiCl_2$ concentrations. **h**, 1D $^{19}$F NMR spectra of isolated FLN5 726His 746His in varying $ZnCl_2$ concentrations. **i,** $^{19}$F NMR spectra of FLN5+34 tfmF655tfmF 726His 746His RNC in varying $NiCl_2$ concentrations, bottom shows Gibb's free energies calculated using lineshape analyses of spectra. Errors were determined by bootstrapping of residuals from lineshape fits. **j,** As in i, but for FLN5+47 with different $^{19}$F-labelled and non-fluorinated variants. **k,** Linewidths determined by lineshape analysis of spectra shown in j. Errors were determined by bootstrapping of residuals from lineshape fits. For all $^{19}$F NMR spectra, observed spectra shown in grey were fitted and assigned to their conformational states (coloured). Total fitted spectra are shown in black, and residual spectrum after fitting are shown below. All spectra were recorded at 500 MHz and 298 K.

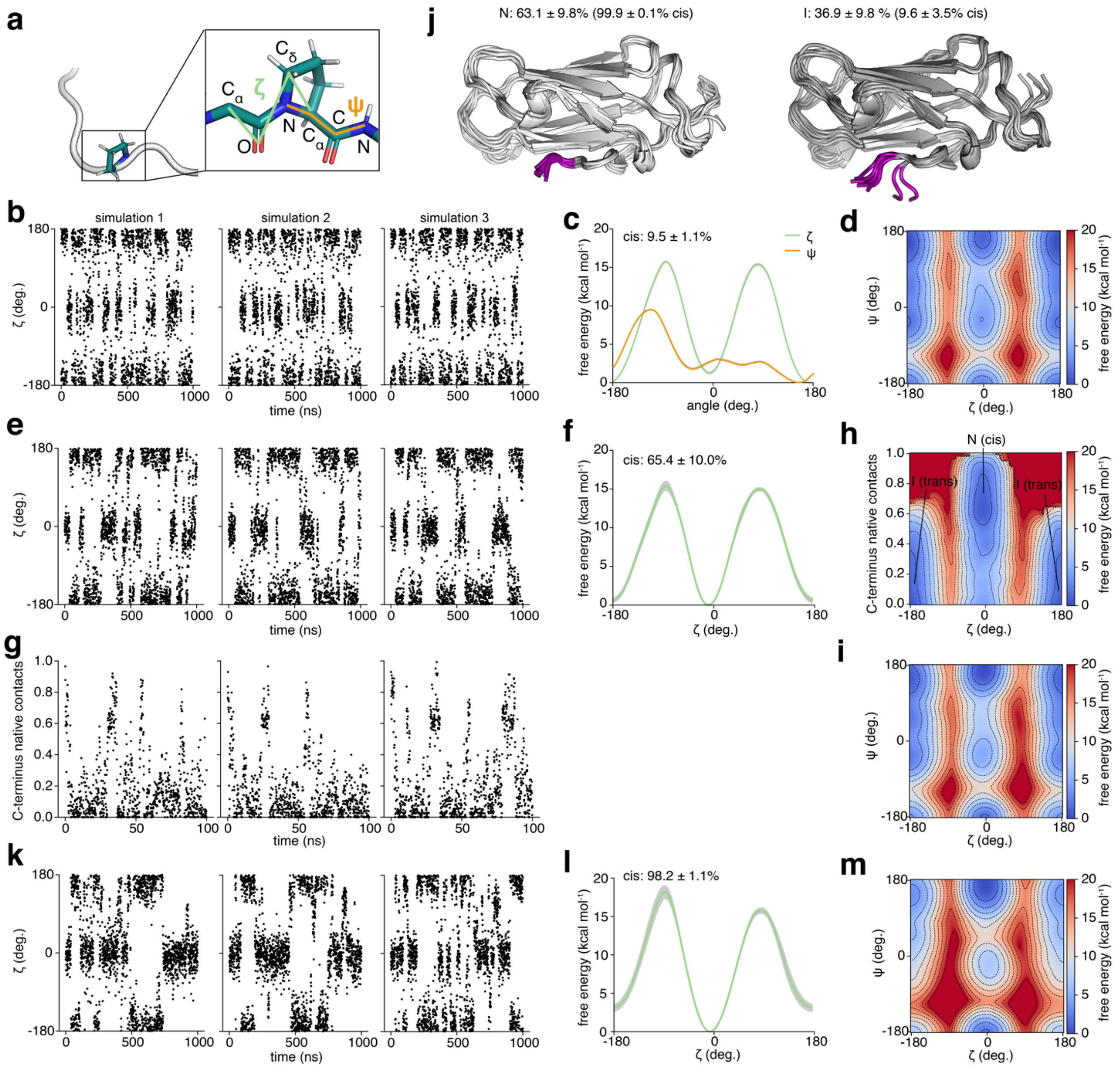


**Figure S4 Structure-dependent proline isomerisation of FLN5 Pro742. a,** Dihedral angles ζ and ψ used as collective variables for enhanced sampling simulations with WT-METAD. **b,** Sampling of ζ in three independent WT-METAD simulations of the unfolded peptide. **c,** Reweighted free energy landscape of Pro742 projected onto ζ and ψ, respectively, for the unfolded peptide (mean ± s.e.m., n=3). **d,** Reweighted 2D free energy landscape of P742 projected onto ζ and ψ for the unfolded peptide. **e,** Sampling of ζ in three independent WT-METAD simulations of FLN5Δ6. **f,** Reweighted free energy landscape of P742 projected onto ζ for FLN5Δ6 (mean ± s.e.m., n=3). **g,** Sampling of C-terminal native contacts (fraction of native contacts formed by residues I743 and D744) in three independent WT-METAD simulations of FLN5Δ6. **h,** Reweighted 2D free energy landscape of FLN5 Δ6 projected onto

$\zeta$ and CTERM averaged across all three simulations. The energy minima corresponding to the native (N) and intermediate (I) state are annotated. **i,** Reweighted 2D free energy landscape of FLN5 $\Delta$6 projected onto $\zeta$ and $\psi$ averaged across all three simulations. **j,** Structural models (10 lowest free energy conformations) corresponding to FLN5$\Delta$6 N and I states and their equilibrium populations predicted by WT-METAD including their P742 cis populations. The C-terminal residues Ile743 and Asp744 are shown in magenta. **k,** Sampling of $\zeta$ in three independent WT-METAD simulations of full-length, folded FLN5. **l.** Reweighted 2D free energy landscape of full-length FLN5 projected onto $\zeta$ and $\psi$ averaged across all three simulations. **m,** Reweighted 2D free energy landscape of full-length FLN5 projected onto $\zeta$ (mean ± s.e.m., n=3).

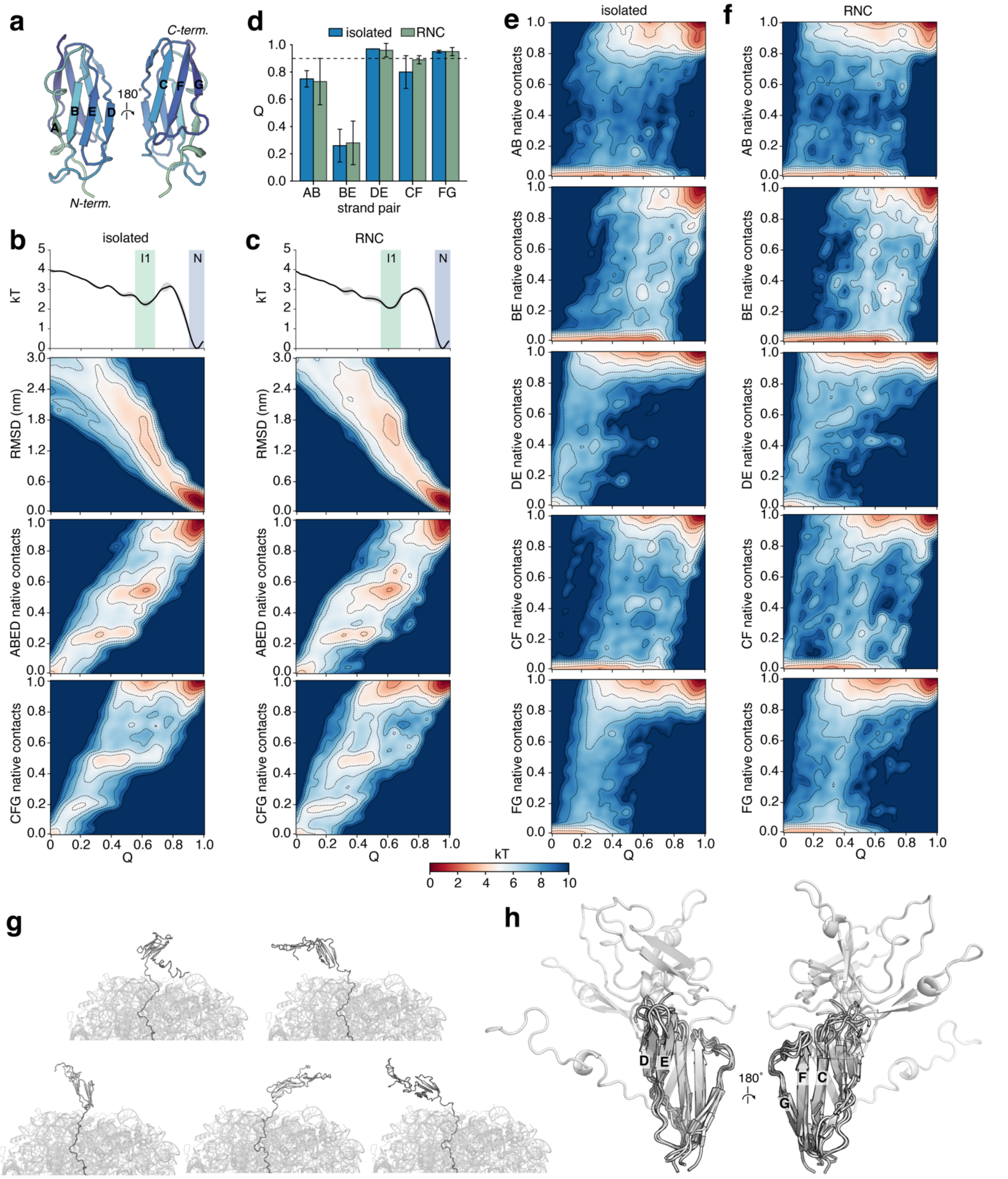


**Figure S5 All-atom folding simulations of FLN5 using rMD. a,** FLN5 crystal structure (PDB 1QFH) annotated with the main β-strands. **b-c,** Kinetic folding free energy landscape of isolated FLN5 (b) and FLN5+47 RNC (c) as a function of the fraction of native contacts (Q), all-atom RMSD with respect to the native structure, and native contacts in sheet 1 (ABED) and 2 (CFG). The 1D energy plot shows the mean and error, estimated as the difference between the first and second half of the dataset. 321 and 226 folding trajectories obtained from rMD simulations were used for the analysis of isolated FLN5 and FLN5+47, respectively. **d,** Average fraction of native contacts calculated for structures attributed to the folding intermediate, I1, energy minimum. Error bars were estimated as the difference between the first and second half of the dataset. **e-f,** Kinetic folding free energy landscape of isolated FLN5 (e) and FLN5+47 RNC (f) projected onto the native contacts of the main β-strand pairs. **g.** Five structural models of I1 on the ribosome, selected for unbiased MD simulations (see Methods). **h.** Aligned structural models of I1 using residues 685-750 (strands C to G) showing the folded core of I1 and disordered N-terminus (transparent, strands A to B).

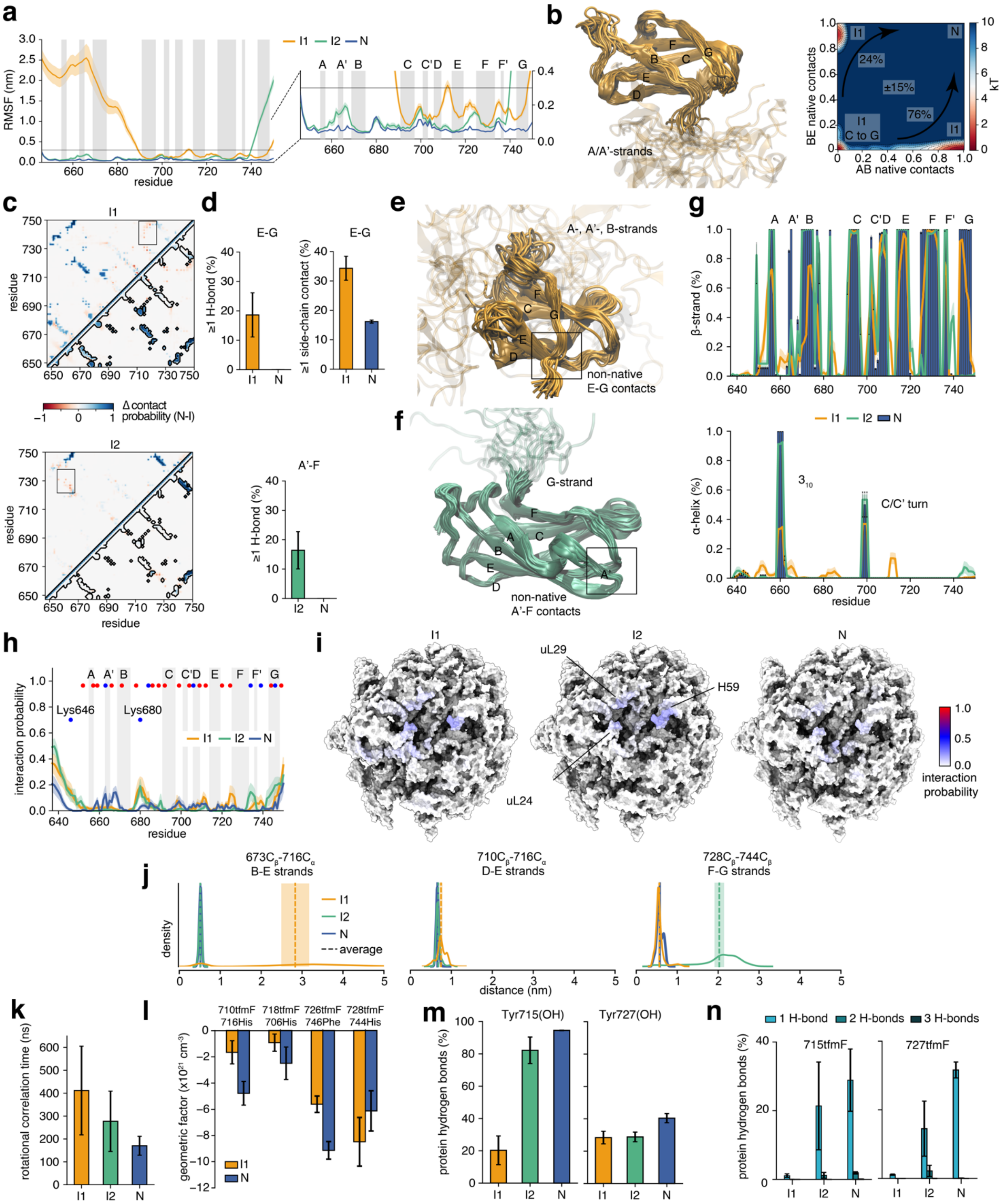


**Figure S6 Structures, dynamics, and interactions of I1 and I2 coTF intermediates. a,** Root mean square fluctuations (RMSF, measured at $C_{\alpha}$ atoms) across the sequence of FLN5 (mean ± s.e.m.) and comparing all conformational states. Secondary structure regions (β-strands) of native FLN5 are annotated. **b,** (left) Structures of I1 from the lowest free energy structural cluster aligned using the common core of strands C to G where the B-strand is also associated with the core. (right) Energy landscape of I1 correlating native contacts between strands A-B (AB) and B-E (BE), showing two parallel folding routes towards the N state. The relative populations of these pathways were estimated from the rMD transition path ensemble (errors represent the difference between the first and second half of the dataset). **c,** Difference contact probability of I1 and I2 compared to N (N-I) highlighting loss of native (blue) and gain of non-native (red) contacts. The black contours represent the native contacts calculated from the crystal structure. Contacts were defined as heavy atom distances of less than 0.5 nm. Two regions in the box highlight non-native contacts (C-G strand contacts in I1 and A'-F strand contacts in I2). **d,** Probability (mean ± s.e.m.) of hydrogen bonds and sidechain contacts (0.35 nm heavy atom cut-off) between non-native structural elements (highlighted in c) compared to the native state. **e,** Representative structures of I1 showing non-native C-G strand contacts. **f,** Representative structures of I2 showing non-native A'-F strand contacts. **g,** Secondary structure propensity of coTF intermediates (lines) compared to the native state shown with bars (mean ± s.e.m.). **h,** Ribosome interaction probability (mean ± s.e.m.) shown along the sequence of FLN5, with native β-strands annotated. Negatively and positively charged residues are denoted with red and blue circles, respectively. **i,** FLN5 (residues 646-750) interactions mapped onto the ribosome surface for all conformational states. **j**, Distance distributions and mean ± s.e.m. of residue sidechains corresponding to $^{19}$F-label pairs calculated for I1, I2 and native state structural ensembles.**k,** Rotational correlation time of FLN5 calculated for all conformational states (mean ± s.e.m.), defined using the natively folded regions for I1 and I2. **l,** Ring current geometric factor ($(1-3\cos^2\theta)/r^3$) (de)shielding contribution to the $^{19}$F chemical shift of probes reporting on strand pair contacts calculated from MD simulations of the native and I1 state using isolated proteins (mean ± s.e.m.). **m,** Tyr-OH hydrogen bonding probability calculated for all conformational state ensembles (mean ± s.e.m.). **n,** Population of hydrogen bonds (H-bonds) between $CF_3$ and protein atoms calculated using isolated protein MD simulations (mean ± s.e.m.).

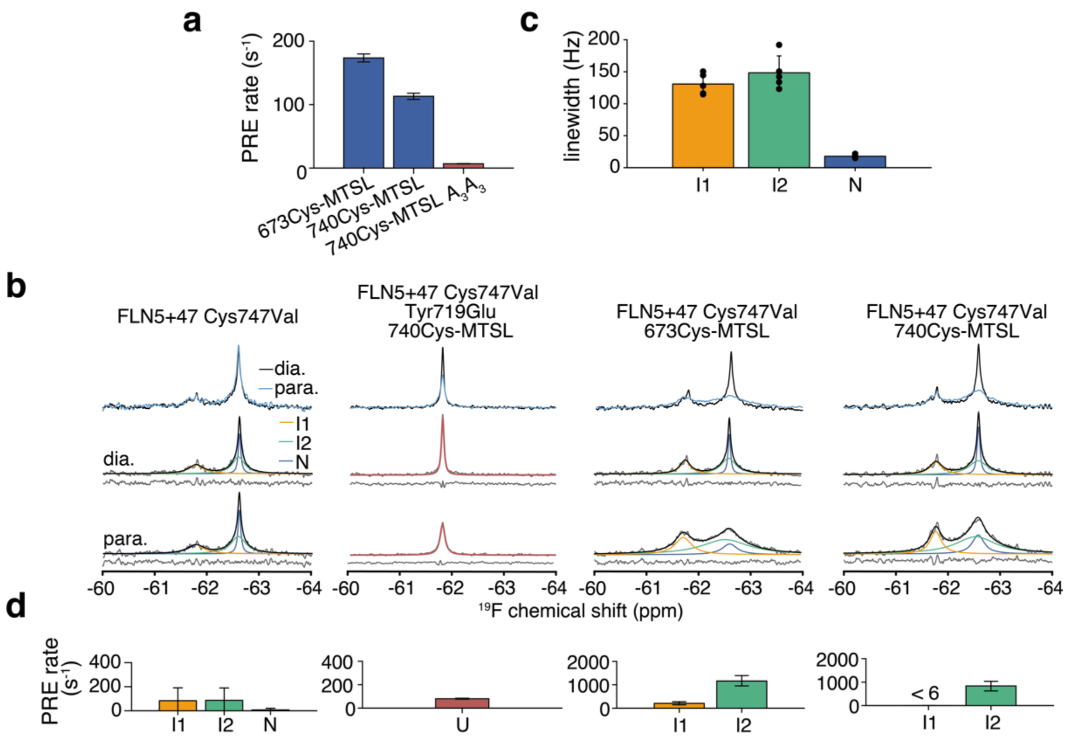


**Figure S7 Validation of structural ensembles using $^{19}$F PRE measurements. a,** PRE rates measured for isolated variants of FLN5 (pseudo wild-type Cys747Val, **Methods**), spectra shown in Figure 4. Errors were determined by bootstrapping of residuals from lineshape fits. **b,** $^{19}$F NMR spectra of FLN5+47 tfmF655 RNC variants under paramagnetic and diamagnetic conditions. Observed spectra shown at top, and also in grey were fitted and assigned to their conformational states (coloured). Total fitted spectra are shown in black, and residual spectrum after fitting are shown below. Spectra were recorded at 500 MHz and 298 K. **c,** Linewidths (mean ± s.d.) measured by lineshape analyses of diamagnetic spectra shown in **b**, used to calculate restraints for lineshape analyses of native state paramagnetic resonances and to determine rotational correlation times for conversion of PRE rates to distances (see **Methods** and **Supplementary Note 4**). **d,** PRE rates calculated from lineshape analyses of spectra shown in **b**. Errors were determined by bootstrapping of residuals from lineshape fits.

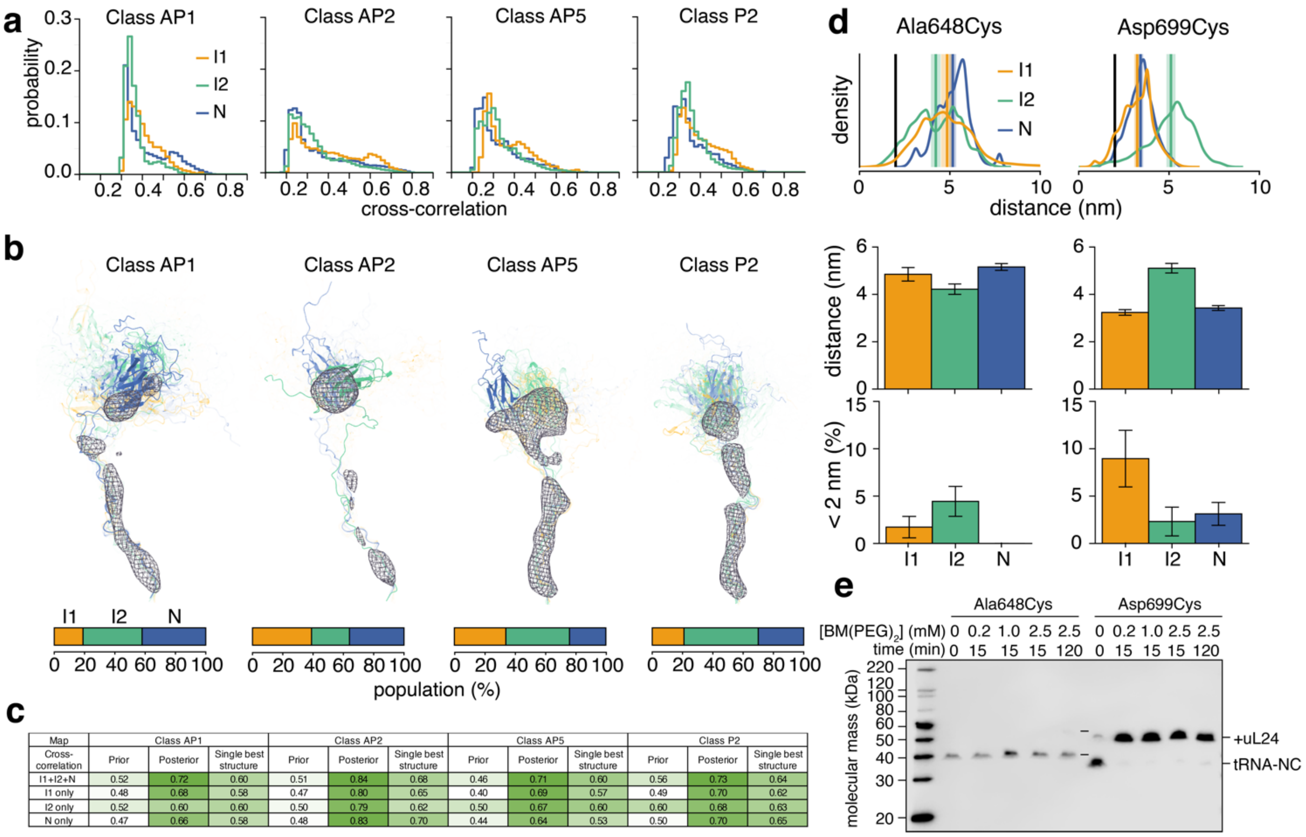


| Map | Class AP1 | | | Class AP2 | | | Class AP5 | | | Class P2 | | |
|---|---|---|---|---|---|---|---|---|---|---|---|---|
| Cross-correlation | Prior | Posterior | Single best structure | Prior | Posterior | Single best structure | Prior | Posterior | Single best structure | Prior | Posterior | Single best structure |
| I1+I2+N | 0.52 | 0.72 | 0.60 | 0.51 | 0.84 | 0.68 | 0.46 | 0.71 | 0.60 | 0.56 | 0.73 | 0.64 |
| I1 only | 0.48 | 0.68 | 0.58 | 0.47 | 0.80 | 0.65 | 0.40 | 0.69 | 0.57 | 0.49 | 0.70 | 0.62 |
| I2 only | 0.52 | 0.60 | 0.60 | 0.50 | 0.79 | 0.62 | 0.50 | 0.67 | 0.60 | 0.60 | 0.68 | 0.63 |
| N only | 0.47 | 0.66 | 0.58 | 0.48 | 0.83 | 0.70 | 0.44 | 0.64 | 0.53 | 0.50 | 0.70 | 0.65 |

**Figure S8 Validation of inter-molecular interactions and orientations of structural ensembles. a-c,** Cryo-EM fitting and reweighting of structural ensembles. **a,** Probability distributions of the correlation coefficients obtained by fitting single conformations from all-atom MD trajectories of the native, I1, and I2 states to one of four cryo-EM maps, corresponding to the AP1, AP2, AP5 and P2 classes of FLN5+47 (**Methods**). **b,** Bayesian reweighting of the collective I1, I2 and N trajectories using the cryoEM maps of FLN5+47, obtained by cryoENsemble[43]. The reweighted structural ensembles are shown, coloured according to their conformational state, and their opacity set to their populations. Total posterior populations of each state are plotted in the bar chart. **c,** Cross-correlations obtained for cryoENsemble reweighting shown in b. **d-e,** Cross-linking experiments of FLN5+47 RNC on uL24 Asn53Cys ribosomes. **d,** Top shows probability distributions and averages (± s.e.m., vertical lines) for all distance pairs (uL24 Asn53 with cysteine residue on FLN5) and conformational states calculated from structural ensembles, middle shows averaged distances (mean ± s.e.m.), and bottom shows probability of contacts using a cut-off of 2 nm. **e,** Anti-hexahistidine detected western blot of FLN5+47 cysteine variant (pseudo wild-type Cys747Val, **Methods**) cross-linked to uL24 Asn53Cys using BM(PEG)$_2$ (**Methods**). Ribosome-bound nascent chain (covalently linked to its tRNA, labelled tRNA-NC) and same but covalently linked to uL24 are indicated.

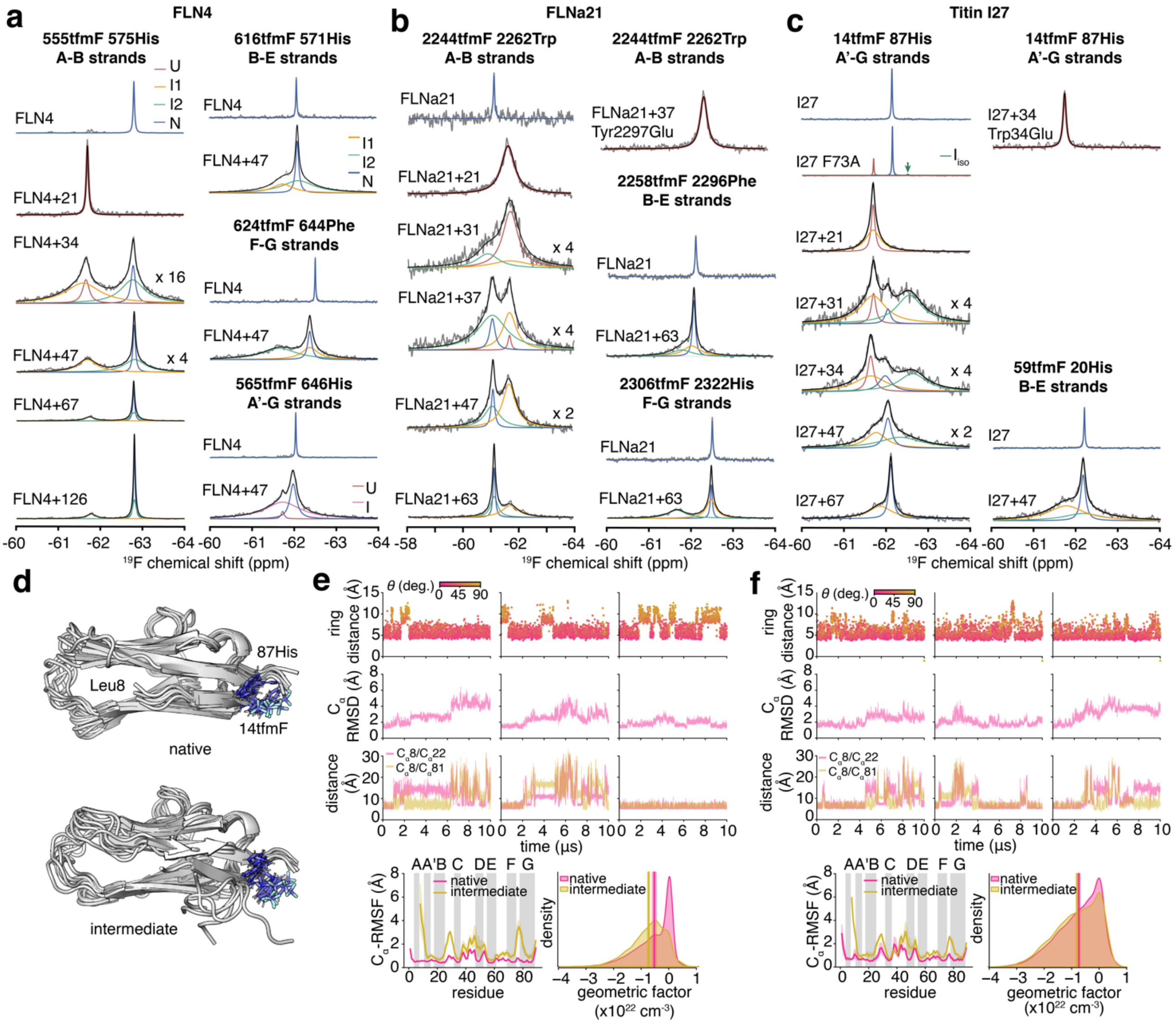


**Figure S9 Characterisation of coTF intermediates of other immunoglobulin-like domains. a,** $^{19}$F NMR spectra of isolated FLN4 and FLN4 RNC variants. **b,** $^{19}$F NMR spectra of isolated FLNa21 and FLNa21 RNC variants. The Tyr2297Glu mutation, analogous to Tyr719Glu in FLN5, was used to destabilise the domain. **c,** $^{19}$F NMR spectra of isolated titin I27 and I27 RNC variants. The Trp34Glu mutation was used to destabilise the domain[3]. Observed spectra shown in grey were fitted and assigned to their conformational states (coloured). Total fitted spectra are shown in black, and residual spectrum after fitting are shown below. Spectra were recorded at 500 MHz and 298 K. **d-g,** Structure, dynamics and aromatic ring currents of the titin I27 folding intermediate (titin I27 ΔA-strand) observed in MD simulations. **d.** Representative structures of the native and intermediate state of titin I27 ΔA. Native was defined as having $C_\alpha$-$C_\alpha$ distances of less than 10 Å for both Leu8-Glu22 and Leu8-Ala81, and intermediate was defined as having at least one of these distances above

10 Å. The sidechains of 14tfmF and 87His are shown in stick representation. **e,** From top to bottom, plot shows centre-of-mass distance between the 14tfmF $CF_3$ and 87His aromatic ring coloured by the angle, θ, formed between the $CF_3$-ring interaction vector and aromatic ring normal vector. Angles below and above 54.7° are considered perpendicular and in-plane $CF_3$-ring interactions. Second plot shows $C_\alpha$-RMSD with respect to the starting structure across all three simulations. Third plot shows distances between the N-terminal $C_\alpha$ (residue Leu8) and $C_\alpha$ atoms of residues Glu22 and Ala81 calculated in three simulations. Bottom left shows average (± s.e.m.) $C_\alpha$ -RMSF of full-length and ΔA titin I27 14tfmF 87His. The full-length simulations were analysed from previous work[39]. The location of β-strands is annotated along the sequence. Bottom right shows geometric factor $(1\text{-}3\cos^2\theta)/r^3$, $r$ is the distance from panel in top left) quantifying the ring current effect experienced by 14tfmF due to 87His for the native and intermediate state. The distribution and mean (± s.e.m. from block averaging) are shown. **f,** Identical analyses as in e for titin I27 ΔA-strand 59tfmF (with wild-type His20).

| Label | RNC | N (Hz) | ± | I2 (Hz) | ± | I1 (Hz) | ± | U (Hz) | ± |
|---|---|---|---|---|---|---|---|---|---|
| 673tfmF | FLN5+47 | 13.9 | 2.8 | 119. | 22.8 | 207. | 39.7 | - | - |
| 710tfmF | FLN5+47 | 25.6 | 2.2 | 344. | 30.6 | 127. | 17.3 | - | - |
| 718tfmF | FLN5+47 | 17.9 | 0.5 | 185. | 10.9 | 184. | 25.2 | - | - |
| | FLN5+47 in urea | 11.3 | 1.3 | 80.3 | 8.7 | 138. | 39.7 | 10.8 | 2.7 |
| | FLN5+34 | 54.9 | 4.0 | 516. | 47.4 | 167. | 42.9 | - | - |
| | FLN5+34 in urea | 55.8 | 3.7 | 549. | 383. | 119. | 27.3 | 21.2 | 2.5 |
| | FLN5+34 | - | - | - | - | - | - | 42.3 | 0.8 |
| 718tfmF | FLN5+47 | 24.2 | 0.7 | 276. | 11.4 | 162. | 15.9 | - | - |
| | FLN5+47 in urea | 26.8 | 4.4 | 85.6 | 22.5 | 152. | 11.5 | 17.6 | 0.5 |
| 726tfmF | FLN5+47 | 16.9 | 0.9 | 281. | 16.9 | 128. | 6.5 | - | - |
| | FLN5+47 in urea | 37.5 | 5.3 | 129. | 34.4 | 148. | 19.5 | 51.5 | 12.0 |
| | FLN5+34 | 57.0 | 3.4 | 255. | 21.7 | 415. | 27.8 | - | - |
| | FLN5+34 in urea | 58.7 | 18.0 | 133. | 10.6 | 219. | 92.9 | 37.1 | 10.8 |
| | FLN5+34 | - | - | - | - | - | - | 184. | 4.5 |
| 728tfmF | FLN5+47 | 25.2 | 1.5 | 218. | 17.3 | 147. | 10.1 | - | - |
| 732tfmF | FLN5+47 | 28.2 | 1.8 | 351. | 23.6 | - | - | 112. | 47.8 |
| | FLN5+47 in urea | 14.9 | 1.4 | 229. | 21.6 | - | - | 79.6 | 5.0 |
| | FLN5+34 | 42.7 | 4.3 | 684. | 132. | - | - | 203. | 56.4 |
| | FLN5+34 in urea | 9.9 | 4.3 | 379. | 30.9 | - | - | 78.1 | 7.9 |
| | FLN5+34 | - | - | - | - | - | - | 267. | 11.1 |
| 715tfmF | FLN5+67 | 66.2 | 2.2 | 365. | 27.8 | 281. | 14.2 | 12.8 | 1.1 |
| 727tfmF | FLN5+67 | 36.0 | 1.0 | 227. | 24.5 | 176. | 14.5 | 21.8 | 11.6 |

**Table S1 Linewidth measurements from lineshape analyses of FLN5 RNCs.** Linewidths determined from lineshape analysis of spectra shown in Figure 1 and Figure S1. Errors were determined by bootstrapping of residuals from lineshape fits.

| Label | RNC | N | ± | I2 | ± | I1 | ± | U | ± |
|---|---|---|---|---|---|---|---|---|---|
| 673tfmF 716His | FLN5+47 | 0.17 | 0.03 | 0.50 | 0.09 | 0.34 | 0.05 | - | - |
| 710tfmF 716His | FLN5+47 | 0.19 | 0.01 | 0.53 | 0.06 | 0.27 | 0.03 | - | - |
| 718tfmF | FLN5+47 | 0.28 | 0.01 | 0.56 | 0.03 | 0.16 | 0.02 | - | - |
| | FLN5+47 in | 0.21 | 0.02 | 0.52 | 0.05 | 0.20 | 0.04 | 0.07 | 0.01 |
| | FLN5+34 | 0.33 | 0.02 | 0.51 | 0.09 | 0.17 | 0.04 | - | - |
| | FLN5+34 in | 0.32 | 0.02 | 0.11 | 0.08 | 0.30 | 0.08 | 0.27 | 0.04 |
| 718tfmF 706His | FLN5+47 | 0.25 | 0.01 | 0.59 | 0.03 | 0.16 | 0.01 | - | - |
| | FLN5+47 in | 0.15 | 0.03 | 0.22 | 0.06 | 0.32 | 0.02 | 0.31 | 0.01 |
| 726tfmF | FLN5+47 | 0.19 | 0.01 | 0.36 | 0.02 | 0.45 | 0.03 | - | - |
| | FLN5+47 in | 0.24 | 0.05 | 0.33 | 0.11 | 0.28 | 0.07 | 0.14 | 0.04 |
| | FLN5+34 | 0.24 | 0.01 | 0.27 | 0.02 | 0.49 | 0.05 | - | - |
| | FLN5+34 in | 0.12 | 0.04 | 0.59 | 0.07 | 0.20 | 0.11 | 0.08 | 0.03 |
| 728tfmF 744His | FLN5+47 | 0.24 | 0.01 | 0.26 | 0.02 | 0.51 | 0.04 | - | - |
| 732tfmF | FLN5+47 | 0.22 | 0.01 | 0.67 | 0.11 | - | - | 0.11 | 0.04 |
| | FLN5+47 in | 0.14 | 0.01 | 0.45 | 0.04 | - | - | 0.42 | 0.02 |
| | FLN5+34 | 0.18 | 0.02 | 0.51 | 0.20 | - | - | 0.31 | 0.10 |
| | FLN5+34 in | 0.03 | 0.01 | 0.61 | 0.07 | - | - | 0.36 | 0.03 |
| 715tfmF | FLN5+67 | 0.34 | 0.01 | 0.23 | 0.02 | 0.31 | 0.02 | 0.12 | 0.01 |
| 727tfmF | FLN5+67 | 0.46 | 0.01 | 0.27 | 0.03 | 0.25 | 0.02 | 0.02 | 0.01 |

**Table S2 Fractional populations from lineshape analyses of FLN5 RNCs.** Populations determined from integrals of lineshapes fitted for spectra shown in Figure 1 and Figure S1. Errors were determined by bootstrapping of residuals from lineshape fits.

| Label | RNC | Time-domain fit BIC | | |
|---|---|---|---|---|
| | | 2-state | 3-state | 4-state |
| 673tfmF 716His | FLN5+47 | 22215 | **22206** | 22228 |
| 710tfmF 716His | FLN5+47 | 24960 | **24901** | 24907 |
| 718tfmF | FLN5+47 | 23313 | **23258** | 23282 |
| | FLN5+47 in | 18665 | 18057 | **17979** |
| | FLN5+34 | 20136 | **20140** | 20160 |
| | FLN5+34 in | 20770 | **17149** | 17161 |
| 718tfmF 706His | FLN5+47 | 23640 | **23427** | * |
| | FLN5+47 in | 22571 | 21910 | **21732** |
| 726tfmF | FLN5+47 | 23912 | **23360** | 23382 |
| | FLN5+47 in | 21906 | 21902 | **21852** |
| | FLN5+34 | 23919 | **23814** | 23815 |
| | FLN5+34 in | 21137 | 21121 | **21080** |
| 728tfmF 744His | FLN5+47 | 23012 | **22722** | 22742 |
| 732tfmF | FLN5+47 | **19339** | 19355 | * |
| | FLN5+47 in | 17263 | **17141** | * |
| | FLN5+34 | **19267** | 19287 | * |
| | FLN5+34 in | 14687 | **14619** | * |
| 715tfmF | FLN5+67 | 30314 | 28851 | **28310** |
| 727tfmF | FLN5+67 | 29544 | 29192 | **29160** |

**Table S3 Time-domain analysis.** Bayesian information criterion (BIC) values calculated for NMR data, displayed in the frequency domain in Figure 1 and Figure S1, analysed in the time-domain using different number of possible resonances, as previously described[2]. The lowest BIC values (in bold) indicate the most likely model. (*) indicates no reasonable fit could be made.

| I1 cluster | Population (%) | I2 Cluster | Population (%) |
|---|---|---|---|
| Cluster_0001 | 22.5 | Cluster_0001 | 25.4 |
| Cluster_0002 | 11.1 | Cluster_0002 | 10.3 |
| Cluster_0003 | 8.8 | Cluster_0003 | 5.3 |
| Cluster_0004 | 5.0 | Cluster_0004 | 4.1 |
| Cluster_0005 | 4.9 | Cluster_0005 | 2.8 |
| Cluster_0006 | 3.7 | Cluster_0006 | 2.4 |
| Cluster_0007 | 3.4 | Cluster_0007 | 2.1 |
| Cluster_0008 | 2.4 | Cluster_0008 | 1.8 |
| Cluster_0009 | 2.2 | Cluster_0009 | 1.7 |
| Cluster_0010 | 2.1 | Cluster_0010 | 1.3 |
| Cluster_0011 | 1.9 | Cluster_0011 | 1.0 |
| Cluster_0012 | 1.7 | Cluster_0012 | 1.0 |
| Cluster_0013 | 1.4 | Cluster_0013 | 0.8 |
| Cluster_0014 | 1.3 | Cluster_0014 | 0.8 |
| Cluster_0015 | 1.2 | Cluster_0015 | 0.7 |
| Cluster_0016 | 1.2 | Cluster_0016 | 0.7 |
| Cluster_0017 | 1.1 | Cluster_0017 | 0.7 |
| Cluster_0018 | 1.0 | Cluster_0018 | 0.7 |
| Cluster_0019 | 1.0 | Cluster_0019 | 0.6 |
| Cluster_0020 | 1.0 | Cluster_0020 | 0.6 |

**Table S4** Populations of top 20 clusters obtained with a RMSD cut-off of 0.2 and 0.1 nm for I1 and I2, respectively.

| Net charge at pH 7.5 (residues) | FLN5 | FLN4 | FLNa21 | I27 |
|---|---|---|---|---|
| Full-length | -9 (K646-G750) | -13 (P547-A648) | -7 (G2236-P2328) | -6 (L1-L89) |
| I1 (C-G strand folded) | -5 (F691-G750) | -8 (F591-A648) | -7 (L2271-P2328) | -1 (V30-L89) |
| I2 (A-F strand folded for filamins, A'-G strand folded for I27) | -10 (K646-L733) | -13 (P547-L631) | -4 (G2236-F2311) | -5 (V11-L89) |

**Table S5** Net charge of proteins studied and their intermediate state folded fragments. Lys/Arg are considered to have a charge of +1, and Asp/Glu have a charge of -1. The favourable ribosome interactions for I2 mediated by FLN5 residues Lys646/Lys680 are structurally conserved in FLN4 (Lys553/Arg583), and FLNa21 (Lys2240/Arg2264).
